# The developmental and genetic architecture of the sexually selected male ornament of swordtails

**DOI:** 10.1101/2020.07.24.219840

**Authors:** Manfred Schartl, Susanne Kneitz, Jenny Ormanns, Cornelia Schmidt, Jennifer L Anderson, Angel Amores, Julian Catchen, Catherine Wilson, Dietmar Geiger, Kang Du, Mateo Garcia-Olazábal, Sudha Sudaram, Christoph Winkler, Rainer Hedrich, Wesley C Warren, Ronald Walter, Axel Meyer, John H Postlethwait

**Affiliations:** Developmental Biochemistry, Biocenter, University of Wuerzburg, Am Hubland, 97074 Wuerzburg, Germany; The Xiphophorus Genetic Stock Center, Department of Chemistry and Biochemistry, Texas State University, San Marcos, Texas, TX 78666, USA; Biochemistry and Cell Biology, Biocenter, University of Wuerzburg, Am Hubland, 97074 Wuerzburg, Germany; Systematic Biology, Department of Organismal Biology, Uppsala University, Norbyvägen 18D, 752 36 Uppsala, Sweden; Institute of Neuroscience, University of Oregon, Eugene, Oregon, OR 97401, USA; Department of Animal Biology, University of Illinois, Urbana, Illlinois, IL 6812, USA; Julius-von-Sachs-Institute for Biosciences, Molecular Plant Physiology and Biophysics, Biocenter, University Würzburg, Julius-von-Sachs-Platz 2, 97082 Würzburg, Germany; Department of Biology, Texas A&M University, College Station, Texas, TX 77843, USA; Department of Biological Sciences and Centre for Bioimaging Sciences, National University of Singapore, Singapore 117543, Singapore; Bond Life Sciences Center, University of Missouri, Columbia, MO USA; Lehrstuhl für Zoologie und Evolutionsbiologie, Department of Biology, University of Konstanz, Universitätsstraße 10, 78457 Konstanz, Germany

## Abstract

Sexual selection results in sex-specific characters like the conspicuously pigmented extension of the ventral tip of the caudal fin - the “sword” - in males of several species of Xiphophorus fishes. To uncover the genetic architecture underlying sword formation and to identify genes that are associated with its development, we characterized the sword transcriptional profile and combined it with genetic mapping approaches. Results showed that the male ornament of swordtails develops from a sexually non-dimorphic prepattern of transcription factors in the caudal fin. Among genes that constitute the exclusive sword transcriptome only two are located in the genomic region associated with this trait; the chaperone, fkbp9, and the potassium channel, kcnh8 that in addition to its neural function performs a role known to affect fin growth. This indicates that during evolution of swordtails a brain gene has been recruited for an additional function in establishing a male ornament.

## Introduction

The evolution of male ornaments has intrigued biologists ever since Charles Darwin struggled to explain how exaggerated, expensive and likely deleterious structures like the peacock’s tail or the horn of male unicorn beetles might have arisen by natural selection. Twelve years after the publication of his book “On the origin of species”, Darwin wrote his second most influential book not on the role of natural, but on sexual selection in evolution [1]. He described the “sword” of the green swordtail, *Xiphophorus hellerii* as an example for his theory on sexual selection and postulated that selection by female choice can be a strong mechanism that could explain the evolution of traits that are clearly detrimental in terms of natural selection [1]. In several species of the genus *Xiphophorus* (Greek for dagger bearer) males carry the sword, a conspicuous extension of the ventral fin rays of the caudal fin that is brightly colored yellow, orange or red and is surrounded by a dark black margin (Fig. 1). The sword develops at puberty and can be as long as the fish itself in some species. Its morphogenesis is instructed by the ventral proximal caudal fin rays, called the “sword organizer” [2]. The sword is a male restricted trait, but female swordtails develop swords like males when treated with testosterone [3, 4]. This suggests that a potential sexual conflict has been solved by a strict androgen dependency for expression of the phenotype. Females of *Xiphophorus hellerii* and several other species preferentially associate with males carrying a longer sword over males with shorter swords, which is thought to result in a higher mating success of long-sworded males [5, 6]. This process exemplifies run-away Fisherian evolution for exaggerated male traits [7]. However, there are also trade-offs [8, 9], because swords attract not only females, but also predators [10], and escape from predators is more difficult because the sword reduces swimming performance [11]. Several species of the genus *Xiphophorus*, including the so-called platyfishes, do not have this sexually dimorphic character (Fig. 1), even though, surprisingly, females nevertheless prefer heterospecific sworded males over their own swordless conspecifics [5]. This observation was used to support a major hypothesis in evolutionary ecology, namely that female preference may drive sexual selection by sensory exploitation since the bias in females was thought to be older than the sword itself [12, 13]. However, molecular phylogenies showed that the sword is an ancestral state [8, 14–16] and implied that derived swordless species had lost the male ornament secondarily, but retained the presumably ancestral female preference for them. This phylogenetic inference fueled the discussion about which evolutionary forces drove the evolution and loss of this conspicuous trait (see [17, 18] [19–21].

**Fig. 1.**
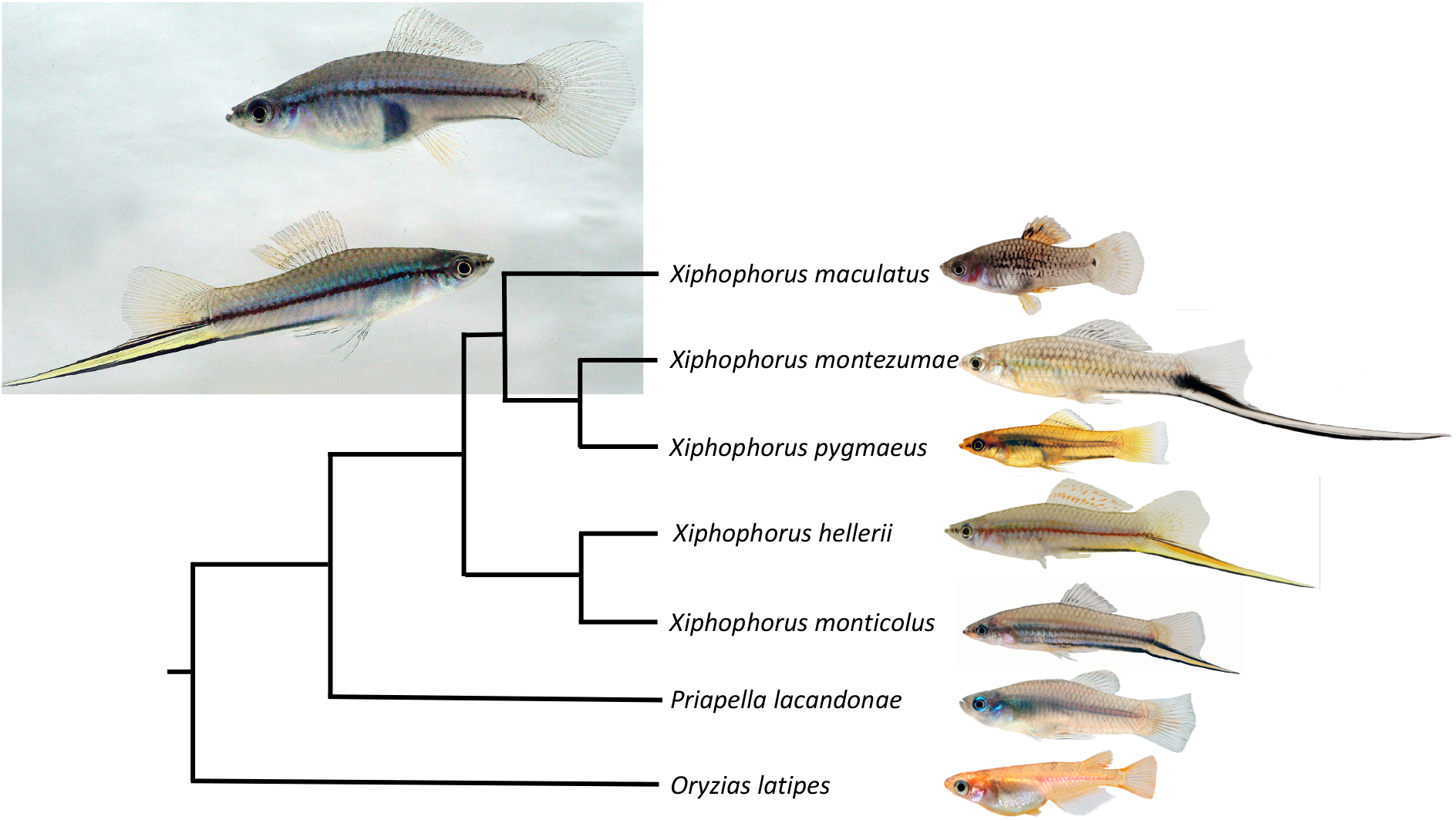
Phylogenetic relationships of sworded and non-sworded *Xiphophorus* species. The swordless *Priapella lacandonae* is the nearest (sister genus) and medaka, *Oryzias latipes*, a distant outgroup. Insert shows female (upper) and male (lower) of the green swordtail, *Xiphophorus hellerii*.

Sword length is a species-specific character and is even polymorphic in two species of Northern swordtails. Females of different *Xiphophorus* species show differences in their preference for sword [5, 22]. Female preferences such as this are considered to potentially not only drive the evolution of male ornaments, but also to result in speciation [23–25]. In the genus *Xiphophorus*, the widespread propensity to prefer sworded males lead to the formation of two hybrid species *X. clemenciae* [8, 21] and *X. monticolus* [16] where, due to the preference for swords females of non-sworded species hybridized with males of swords species to bring about new, sworded hybrid species.

A huge body of literature on how both sexual and natural selection can lead to speciation has been published[26, 27] but almost nothing is known about the genetic basis of male ornaments or male “weapons” used in male-male competition [28, 29]. To identify the genes on which female preferences act on is an important task that is necessary to permit the testing of hypotheses regarding the roles of sexual selection at the molecular genetic level.

The swords of swordtails became a textbook example of a sexually selected trait, yet despite research efforts for almost three decades the molecular genetic basis of sword development remained unkown. So far, candidate gene approaches involving known genes of fish fin growth and development [30] [31] and suppression subtractive hybridization cloning [32] have not revealed the secret of the sword.

To identify the genetic basis for sword formation, we combined genome-wide expression analysis during sword development and regeneration with a genetic association study for sword length in a cross of a non-sworded species to a sworded species.

## Results

To obtain a most comprehensive list of protein coding genes that are involved in the formation of the sword, we compared expression levels using several RNA-seq datasets from the green swordtail, *Xiphophorus hellerii* (Fig. 1). We reasoned that sword genes should be differentially expressed (i) during growth of the developing sword of males at puberty (fig. S1) and (ii) during the course of sword regeneration (fig. S2). Because immature fish and adult females also develop a sword indistinguishable from the male structure following treatment with androgens [3, 4] we generated (iii) one RNA-seq dataset from testosterone-treated adult females; and added (iv) our previous dataset from testosterone-induced swords in pre-pubertal juveniles [3]. Small biopsies from the dorsal and ventral fin margin during a timed series of growth and of regeneration and from the hormone induced and naturally developed swords from 15-20 individuals were pooled and used for transcriptome sequencing. Differential expression was deduced from comparison to the corresponding dorsal part of the caudal fin. The four datasets were overlapped to identify genes that are commonly regulated in all four processes of sword development (fig. S3). This process yielded a set of 68 regulated genes differentially expressed (log2FC >=1) in all sword transcriptomes (11 down and 57 upregulated, table S1).

We expected differentially expressed genes to be of two main categories: those primarily responsible for inducing the sword and those that execute the instruction process by actually building the components of the sword. The sword, like other parts of the caudal fin, consists of bony fin rays, skin, pigment cells, sensory neurons, blood vessels and mesenchyme. Amongst genes upregulated in sword vs control fin regions, four genes (*xdh, tyr, myrip, agrp*) are obviously connected to sword pigmentation; several other upregulated genes are related to increased vascularization (*angptl5, angptl 1)* and fin-ray rigidity (collagen*s col9a1, col10a1* and extracellular matrix proteins *fib7l, spock 2, tn-c, frem3, cd200, and4, gpc2*) that support the sword structure as an extremely long outgrowth of ventral fin rays. It is unclear whether these genes are also critical for the primary process of induction and development of the sword, but all are reasonably predicted to be involved in later differentiation processes. The sword transcriptome was also enriched for genes with neural functions (*pdyn*, *draxin*, *kcnh8*, *kcng2*, *chrna7*, *ncan*, *nrxn*, *lypd6, gfra1*) and Ca^2+^ signaling (*stc2*, *efcc1*, *fkbp9*, *fkbp11*).

Intriguingly, several transcription factors were included in the differentially expressed genes list and could be strong candidates for having a critical function in regulating caudal fin development and consequently also sword formation. *Homeobox protein six2a*, which plays a role in chicken hindlimb development [33], forms a continuous dorsoventral expression gradient in the swordtail tail fin (Fig. 2A, table S2), similar to several developmental transcriptional regulators in the establishment of the zebrafish pectoral fin anterior-posterior axis [34]. The dorsalizing factor *zinc finger protein zic1*, which is critical for the development of the homocercal fin shape in fish [35] is highly expressed in the dorsal compartment, but expression is absent from the medial region and all sword transcriptomes (table S2). More strikingly, *homeobox protein hoxb13a*, which is the most caudally expressed *hox* gene in fish [36], has high expression in the non-sword regions of the *X. hellerii* caudal fin, but is not expressed in the sword and the sword-organizer (table S2). During tail fin regeneration, *hoxb13a* is upregulated in the median and dorsal region but not expressed in the outgrowth leading to the sword (Fig. 2). The *t-box transcription factor tbx3a* gene, which promotes formation of the mesoderm cell lineage [37] and is involved in vertebrate limb pattern formation [38], is lowly expressed in the non-sword regions of the tail fin, but abundant in the sword organizer region at the base of the fin, and in the sword during regeneration, natural sword development and hormone-induced sword (Fig. 2, table S2). The same expression pattern is displayed by *paired box protein pax9*, which in fish is a critical factor for development of the hypural plate supporting the peduncle [39], where the caudal fin is inserted (Fig. 2, table S2). Interestingly, *leukocyte tyrosine kinase receptor* (*ltk*), which normally has no spatial expression pattern in the caudal fin of *X. hellerii* males, builds up a local expression pattern in the sword producing blastema similar to that of *hoxb13a* during caudal fin regeneration and natural and hormone induced sword development (fig. S4, table S2).

**Fig. 2.**
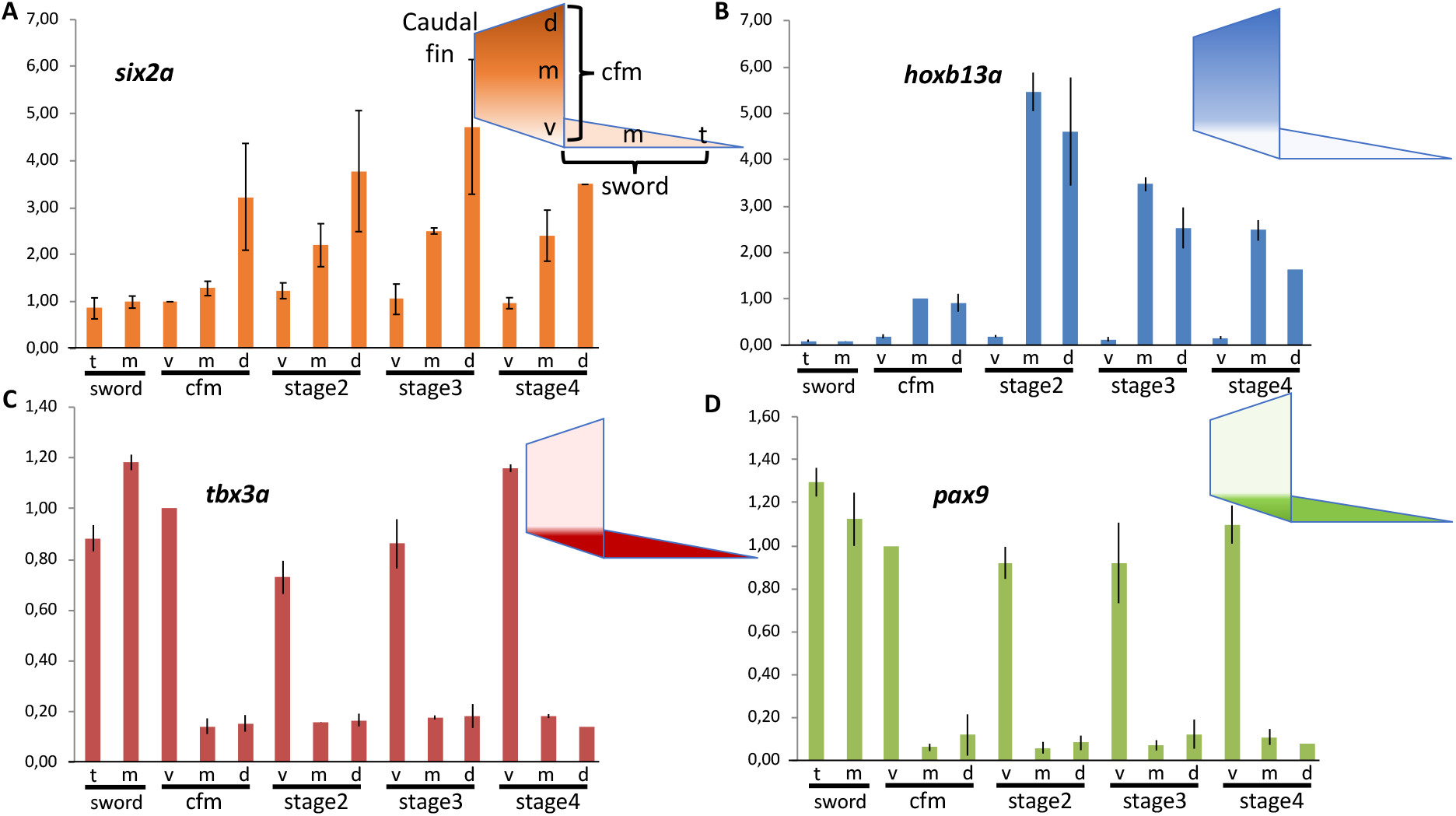
Spatial expression pattern of transcription factor genes in the caudal fin and sword of male *Xiphophorus hellerii*. Expression of *six2a* (A), *hoxb13a* (B), *tbx3a* (C) and *pax9* (D) in the caudal fin margin of the tail fin (cfm) of adult *Xiphophorus hellerii* males, the median sector (m) and tip (t) of the sword and during sword regeneration (v, ventral, m, median, d, dorsal compartment). Vertical axis indicates fold change of expression normalized to cfm, v (*six2a, tbx3a, pax9*) or cfm, m (*hoxb13a*).

Males of two other swordtail species, *X. montezumae* and *X. monticolus* (fig. S5, 6) showed the same expression gradients and temporal pattern during sword regeneration. Of note, analysis in *X. montezumae*, the species with the longest sword (sword index = sword length/standard body length up to 1.9), revealed that the transcription factor expression pattern is immediately initiated in the blastema of the regenerating caudal fin and builds up to the levels of the caudal fin margin and sword during the first days of growth. The platyfish *X. maculatus*, a species which does not develop a sword, and the pygmy swordtail, *X. pygmaeus*, where males have only a tiny unpigmented ventral protrusion of the tail fin but no sword, display the transcription factor gradients in the caudal fin, but these gradients are much less pronounced and at lower transcript levels (fig. S7-9). Phylogenetic evidence suggested that these species have lost the sword secondarily [8, 14]. Apparently, the loss of the male ornamental trait is associated with a decay of this gene expression pre-pattern. The sword arose at the basis of the genus *Xiphophorus* [8, 14]. In, *Priapella*, a swordless sister genus, the tail fin pattern on which the sword is built is already present to a large extent. The expression patterns of *pax9*, *tbx3* and *six2a* are conserved, only *hoxb13a* expression is in additional absent from the dorsal compartment (fig. S8, 9). In the distantly related medaka, *Oryzias latipes*, the tail fin spatial expression patterns of *hoxb13* and *pax9* are like in *Xiphophorus*, however, at much lower transcript levels. However, expression of the medaka orthologs of *tbx3* and *six2a* is not detected in the caudal fin (fig. S9).

Importantly, the same expression profile for all five transcription factors was also observed in female swordtail caudal fins (fig. S10, table S1, S2), although at lower expression levels for *six2a*, *tbx3a* and *pax9*. However, this finding indicates that a pre-pattern of transcription factors exists in the caudal fin of both sexes that provides in males the positional information for sword development, but this rules out those genes as candidates for sword induction.

Reasoning that genes that are responsible for sword would be expressed only in males, we thus generated transcriptomes from upper and lower terminal caudal fin compartments of females and used these to eliminate genes from candidate status in the sword transcriptome if they showed the same regulation in male and female caudal fin regeneration. This process still left us with 54 candidate genes (table S1). To further reduce the number of genes we performed a genetic mapping approach.

Thus, we performed QTL mapping using RAD-tags. Because crossing of a swordtail to a nearest outgroup species prior to evolution of this character (e.g. *Priapella sp.*) is not possible, we used a congeneric species that has lost the sword. A backcross between the sword-less Southern platyfish *X. maculatus* and the green swordtail *X.hellerii* using *X.hellerii* as the recurrent parent was generated [40]. Mapping the sword-index of 85 backcross males against genetic polymorphisms in the reference swordtail genome revealed significant association with a region on linkage group (LG) 13 (LOD score max likelyhood = 3.86, non-parametric = 4.87) (Fig. 3, fig. S11). A region on LG 1 (LOD score ml = 3.17, np = 1.57) and LG 9 (LOD score ml = 2.54, np =2.15) barely failed to reach the significance level.

**Fig. 3.**
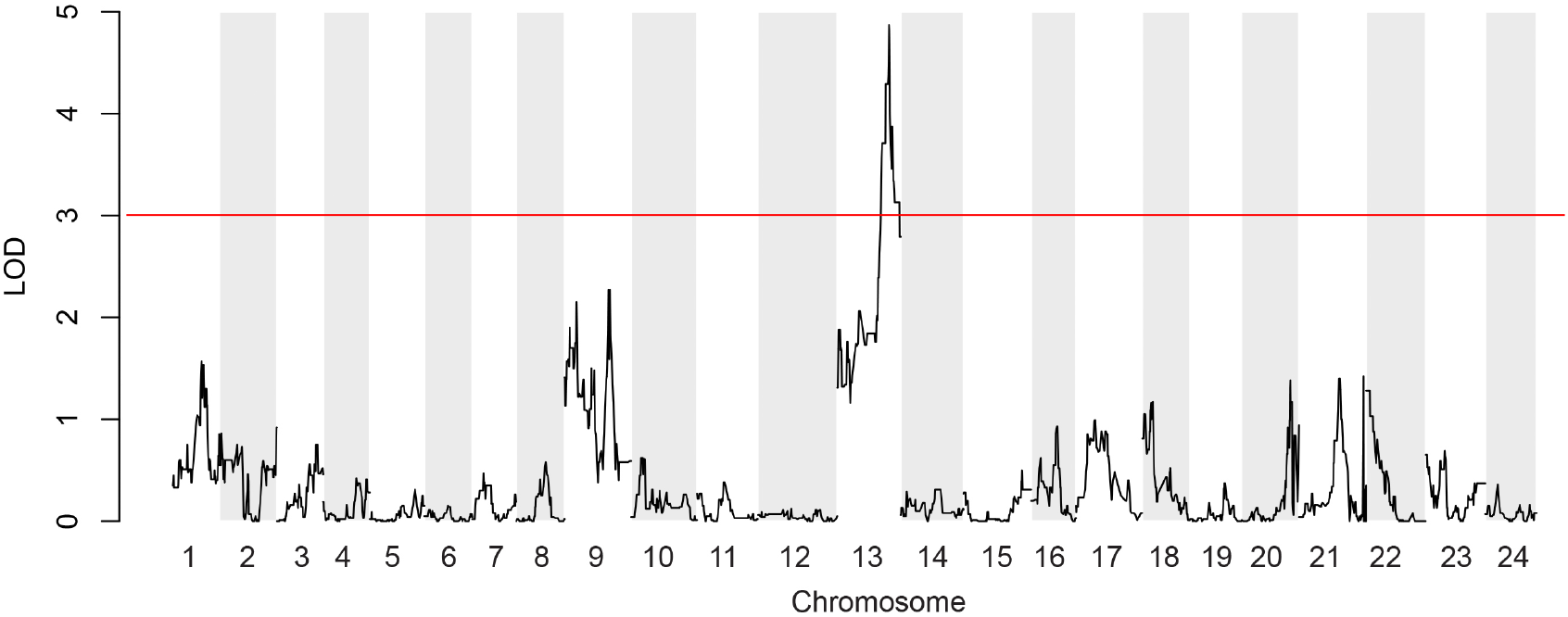
Manhattan plot of quantitative trait loci (QTL) mapping results for sword length. One major QTL peak is located on chromosome 13, two minor peaks on chromosomes 1 and 9. The plot depicts aligned RAD-tag positions on the *Xiphophorus hellerii* genome version 4.1 with non-parametric statistics

Several minor peaks also appeared on LG’s 20 – 24. This result defines the sword as a highly polygenic trait, which is in accordance with the size distribution of sword lengths in platyfish/swordtail hybrids [41].

When the positions of sword specific differentially expressed genes (table S1) were examined with respect to the QTL peaks in the 2.0 LOD interval, none of the genes involved in establishing the prepattern and none of the pigmentation, angiogenesis, or ECM genes that were differentially regulated during sword development were found to be encoded in any of the regions identified in the QTL analysis. Only two differentially expressed genes with log2FC >=1 mapped to a QTL peak, both in the main peak on chromosome 13. These are *fkbp9* and *kcnh8*.

The gene encoding the chaperone peptidyl-prolyl cis-trans isomerase Fkbp9 is 2-to 3-fold higher expressed in the developing sword than in control tissue and becomes upregulated in sword regeneration at stages 3-4 (fig. S12, table S2). Expression is not elevated in the sword organizer, which weakens its candidacy as a gene responsible for induction of sword development.

The other gene that has overlapping candidacy from both gene expression and mapping studies is *kcnh8.* Kcnh8 is a potassium channel of the *ether-à-go-go* (EAG) type that is expressed abundantly in brain and at intermediate levels in ovary and testis (Fig. 4A). Importantly, *kcnh8* is strongly upregulated in the sword during normal development and following androgen treatments, in the sword organizer region, and in the fully developed sword, and becomes strongly upregulated during sword regeneration (Fig. 4B, table S2). It is always amongst the 0.3% of most differentially expressed genes (>21,000 total). Transcripts of *kcnh8* are almost absent from all other fin areas of males and *kcnh8* is only expressed at background levels in female caudal fins.

**Fig. 4.**
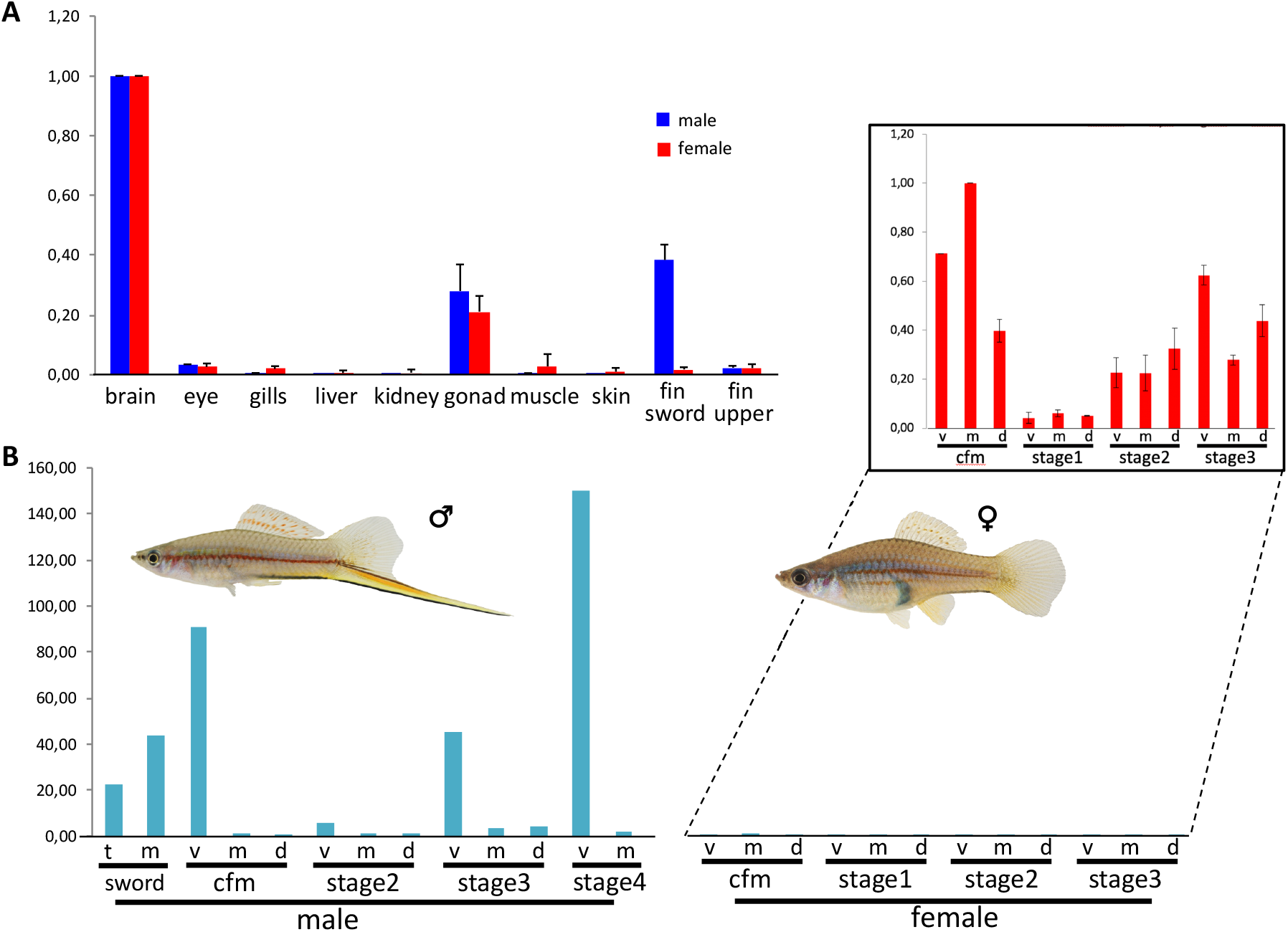
Expression of *kcnh8* in adult males and females of Xiphophorus hellerii. (A) Organ-specific expression profile in adult females and males. (B) Expression of *kcnh8* in the caudal fin margin of the tail fin (cfm) of adult *Xiphophorus hellerii* males and females, the median sector (m) and tip (t) of the sword and during caudal fin regeneration (v, ventral, m, median, d, dorsal compartment). Insert: expression in females upscaled. Vertical axis indicates fold change of expression normalized to brain (A), cfm, m (B).

Expression of swordtail Kcnh8 in the Xenopus oocyte system and two-electrode voltage clamp analyses revealed that the protein has the hallmark characteristics of a fully functional voltage gated potassium channel member of the K_v_12.1 family[42] in terms of voltage activation characteristics, time-dependent activation kinetics, potassium selectivity and inhibition by Ba^2+^ ions (Fig. 5).

**Fig. 5.**
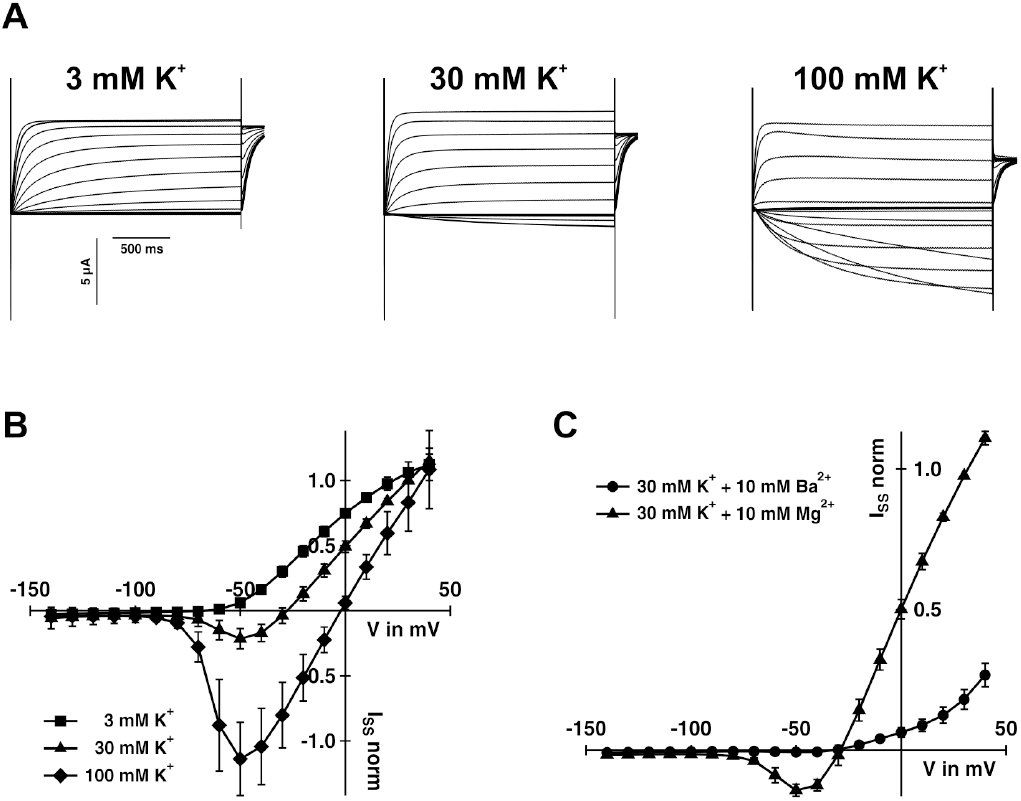
Electrical features of *Xiphophorus hellerii* Kcnh8. (A) Representative TEVC recordings of KCNH8-expressing Xenopus oocytes at the indicated potassium concentrations. Test voltages ranged between +40 to −140 mV in 10 mV decrements. (B) Steady-state currents (I_SS_) extracted from recordings as shown in A) of KCNH8-expressing oocytes were plotted as a function of the applied membrane potential (mean of n ≥ 7 oocytes ± SD of ≥ 3 independent experiments). (C) Application of 10 mM BaCl_2_ in the presence of 30 mM KCl inhibited the KCNH8-mediated I_SS_ (mean of n = 6 oocytes ± SD of ≥ 2 independent experiments). (B) and (C) I_SS_ were normalized to the currents at +30 mV in standard bath medium (30 mM KCl).

We found that also *X. montezumae*, which has an even longer sword than *X. hellerii*, has the same high expression of *kcnh8* in the sword and during sword regeneration (fig. S13). Interestingly, in species that develops shorter sword than *X.hellerii* or only tiny protrusions swords, *X. monticolus* and *X. pygmaeus*, *kcnh8* expression during sword regeneration is only weakly upregulated. In the swordless platyfish *X. maculatus*, no differential expression of *kcnh8* was noted between the lower and upper compartment and during regeneration of the caudal fin (fig. S13).

## Discussion

Sexually selected traits are present in many species and a hallmark of sexual dimorphism between males and females. The evolutionary mechanism driving their origin, maintenance and role in speciation have been widely studied, but today little is known about the proximate causes, i.e. the genes encoding sexually selected traits and their function in development of the structure, aside a few examples from Drosophila [43, 44]. The sword is a male specific outgrowth of the lower margin of the caudal fin and we wanted to know what genes provoke its sex-specific elongation. The fins of fish are intricate three-dimensional structures composed of numerous cell types. Size, shape, pigmentation and other features of fins are generally highly fixed and specific for different species and certain ontogenetic stages. In many species fins are sexually dimorphic traits [45]. In zebrafish it has been shown that pectoral fins have a regionalized gene expression pattern that creates gradients of transcription factors [34]. We conclude that also in the caudal fin of male swordtails a similar specific regionalized gene activity pattern provides the positional information for development of the sword. The regional expression of the transcription factors Hoxb13a, Six2a, Tbx3a and Pax9 produces a prepattern in the tail fin that is connected to sword development since the expression pattern vanishes in species that have secondarily lost the sword. This pattern is established before the sword develops during puberty and its presence (with minor deviations) in adult females may allow the development of a sword after experimental androgen treatment or as a natural phenomenon in old post-reproductive females [46, 47].

To identify those genes that are determining the development of the sword in males we reasoned that such genes should be differentially expressed in sword development and encoded in genomic regions that are linked to this trait. Our QTL analysis, consistent with earlier genetic findings [41], uncovered that several chromosomal regions contribute to the polygenic basis of the male structure. Consistently, the major locus on chromosome 13 fully overlaps a similar broad QTL that was obtained in an independent study for the character sword length in natural hybrids between a swordless (*X. birchmanni*) and a sworded (*X. malinche*) Northern swordtail species[48].We identified two candidate genes that appear to be involved in the development of the sword. Rather than being typical regulators of development and differentiation such as transcription factors or extracellular diffusible growth factors, experiments identified a channel protein, *kcnh8*, and a chaperone, *fkpb9*.

In zebrafish long fin mutants, mutations in several potassium channel genes, including *kcnh2a*, *kcnk5b*, and *kcc4a* cause various types of fin overgrowth [49–51]. In fighting fish, *Betta splendens, kcnh8* mis-expression is associated with pectoral fin overgrowth (Wang et al. submitted). A hyperpolarizing mutation in *kcnk5b* causes the long fin phenotype in ornamental goldfish [52]. Mutations disrupting ion channels and ion-dependent signaling are extensively related to abnormal organ development and regeneration via bioelectrical regulation [53]. Potassium channels of the Kcnh family have been implicated in cell proliferation by influencing membrane polarization and thus calcium signaling [54, 55]. Increased intracellular calcium levels activate osteoblasts and their precursors [56, 57], which build the fin rays of the overgrowing structures of the long-fin mutants and the *Xiphophorus* sword. Potassium channels can also play a role in cell cycle and proliferation control by mechanisms unrelated to ion channel permeability [55]. Despite this wide spectrum of biological functions of potassium channels besides the classical channel properties, their transcriptional regulation and biochemical interactions are not well understood.

Voltage gated channels of the EAG family are inhibited by intracellular calcium [58]. One function of Fkpb9 besides acting as a prolyl cis-trans isomerase is mediated through its calcium binding Ef-H domain [59]. In zebrafish tailfin growth a predominant role for the calcium activated protein phosphatase calcineurin was shown. In this case inhibition of this pathway led to unscheduled outgrowth of the caudal fin margin [60].

Kcnh8 is the pore forming unit of some voltage-gated potassium channels, which have broad functions mainly in neurotransmitter release and neuronal excitability, but also in epithelial electrolyte transport and cell volume regulation [55, 61]. In zebrafish, due to the presence of duplicate versions of the channel protein coding genes, one paralog obviously can fulfill functions restricted to the fin. Mutations of the “fin” paralog only affect fin growth, while the other channel functions are executed by the second paralog. However, *kcnh8* is present only as a single copy and it is abundantly expressed in the brain and to a lesser extent in the gonads of both sexes and additionally only in the male sword of *Xiphophorus* but importantly not in the corresponding part of the female caudal fin. These expression domains imply that a neuronal gene was recruited during the evolution of the male ornament about 3-5 million years ago, early during the diversification of swordtail fish through a rewiring of its regulatory network rather than by selection on its protein function. The Kcnh8 proteins of *Xiphophorus* species have a few aminoacid changes, which, however, do not correlate with the presence or absence of a sword in males (fig. S14). Thus, it is more likely that the function for sword development has been added to the *kcnh8* gene through changes in gene regulation.

The implication of Kcnh8 activity in natural sword development adds a case of an evolutionary mutant for a potassium channel being involved in regulation of fin growth, which thus far were only seen in laboratory mutants. It appears that the four genes, *kcnh2a*, *kcnk5b*, *kcc4b* and *kcnh8,* govern a common pathway of downstream signaling that connects membrane potential, K+ permeability, eilennummern and calcium homeostasis to the ubiquitous machinery of cell growth and proliferation. Although swordtails, because of their livebearing mode of reproduction are not amenable to transgenic technologies, the induced fin mutants of egg laying fish can be employed to systematically knock-out candidate signal transducers and elucidate the interface between ion channels and growth control.

## Acknowledgements

This work was supported by the Deutsche Forschungsgemeinschaft (grants 5263398, 163418330 and 5446040 to AM) and by NIH grant 5R01OD011116 (JHP), 5R24OD018555 (JHP, MS, RW, WW).

## Authors contributions

MS, AM and JHP conceived the study and coordinated the work. JA, AA, JC, JW and JHP did the QTL mapping, JO and CS prepared RNA and performed the qRT-PCR experiments, DG and RH characterized the channel properties of *Xiphophorus* Kcnh8, SS and CW analyzed sword growth and regeneration, SK, DK and MGO analyzed the RNA-seq data and intersected the expression with the QTL data, AM contributed RNA-seq data from androgen induced swords, WCW and RW contributed the *Xiphophorus hellerii* genome, MS analyzed all data and drafted the manuscript, all authors were involved in preparing the final version of the manuscript.

## References

1. Darwin, C., The Descent of Man and Selection in Relation to Sex. 1871, London: John Murray.

2. Eibner, C., et al., An organizer controls the development of the “sword,” a sexually selected trait in swordtail fish. Evol Dev, 2008. 10(4): p. 403–12.

3. Kang, J.H., et al., Transcriptomics of two evolutionary novelties: how to make a sperm-transfer organ out of an anal fin and a sexually selected “sword” out of a caudal fin. Ecol Evol, 2015. 5(4): p. 848–64.

4. Dzwillo, M., Sekundäre Geschlechtsmerkmale an der Caudalflosse einiger Xiphophorini unter dem Einfluß von Methyltestosteron. Mitt. Hamburg. Zool. Mus. Inst, 1964(61): p. 15–22.

5. Basolo, A.L., Female preference predates the evolution of the sword in swordtail fish. Science, 1990. 250(4982): p. 808–10.

6. Basolo, A.L., Evolutionary change in a receiver bias: a comparison of female preference functions. Proc Biol Sci, 1998. 265(1411): p. 2223–8.

7. Fisher, R.A., The Genetical Theory of Natural Selection. 1930, Oxford: Oxford University Press.

8. Meyer, A., J.M. Morrissey, and M. Schartl, Recurrent origin of a sexually selected trait in Xiphophorus fishes inferred from a molecular phylogeny. Nature, 1994. 368(6471): p. 539–42.

9. Wagner, W.E., Jr., et al., Tradeoffs limit the evolution of male traits that are attractive to females. Proc Biol Sci, 2012. 279(1739): p. 2899–906.

10. Rosenthal, G.G., et al., Shared preferences by predators and females for male ornaments in swordtails. Am Nat, 2001. 158(2): p. 146–54.

11. Basolo, A.L. and G. Alcaraz, The turn of the sword: length increases male swimming costs in swordtails. Proc Biol Sci, 2003. 270(1524): p. 1631–6.

12. Endler, J.A., Signals, signal conditions, and the direction of evolution. American Naturalist, 1992. 139: p. 125–153.

13. Ryan, M.J., Sexual selection, sensory systems and sensory exploitation. Oxford Surveys in Evolutionary Biology, 1990. 7: p. 157–195.

14. Jones, J.C., et al., The evolutionary history of Xiphophorus fish and their sexually selected sword: a genome-wide approach using restriction site-associated DNA sequencing. Mol Ecol, 2013. 22(11): p. 2986–3001.

15. Cui, R., et al., Phylogenomics reveals extensive reticulate evolution in Xiphophorus fishes. Evolution, 2013. 67(8): p. 2166–79.

16. Kang, J.H., et al., Comprehensive phylogenetic analysis of all species of swordtails and platies (Pisces: Genus Xiphophorus) uncovers a hybrid origin of a swordtail fish, Xiphophorus monticolus, and demonstrates that the sexually selected sword originated in the ancestral lineage of the genus, but was lost again secondarily. BMC Evol Biol, 2013. 13: p. 25.

17. Pomiankowski, A., Sexual selection. Swordplay and sensory bias. Nature, 1994. 368(6471): p. 494–5.

18. Da Silva, J., Male swords and female preferences. Science, 1991. 253(5026): p. 1426.

19. Winquist, S.T., et al., Male swords and female preferences. Science, 1991. 253(5026): p. 1426.

20. Endler, J.A. and A.L. Basolo, Sensory ecology, receiver biases and sexual selection. Trends Ecol Evol, 1998. 13(10): p. 415–20.

21. Meyer, A., W. Salzburger, and M. Schartl, Hybrid origin of a swordtail species (Teleostei: Xiphophorus clemenciae) driven by sexual selection. Mol Ecol, 2006. 15(3): p. 721–30.

22. Wong, B.B. and G.G. Rosenthal, Female disdain for swords in a swordtail fish. Am Nat, 2006. 167(1): p. 136–40.

23. Coyne, J.A. and H.A. Orr, Speciation. 2004, Sunderland, MA: Sinauer.

24. Mayr, E., Animal species and evolution. 1963, Cambridge, MA: Belknap Press of Harvard University Press.

25. Ritchie, M.G., Sexual selection and speciation. Annual Review of Ecology, Evolution, and Systematics, 2007. 38: p. 79–102.

26. van Doorn, G.S., P. Edelaar, and F.J. Weissing, On the origin of species by natural and sexual selection. Science, 2009. 326(5960): p. 1704–7.

27. Tinghitella, R.M., et al., On the role of male competition in speciation: a review and research agenda. Behavioral Ecology, 2017. 29(4): p. 783–797.

28. Gotoh, H., et al., The function of appendage patterning genes in mandible development of the sexually dimorphic stag beetle. Dev Biol, 2017. 422(1): p. 24–32.

29. Kraaijeveld, K., Genetic architecture of novel ornamental traits and the establishment of sexual dimorphism: insights from domestic birds. Journal of Ornithology, 2019. 160(3): p. 861–868.

30. Zauner, H., et al., Differential regulation of msx genes in the development of the gonopodium, an intromittent organ, and of the “sword,” a sexually selected trait of swordtail fishes (Xiphophorus). Evol Dev, 2003. 5(5): p. 466–77.

31. Offen, N., et al., Fgfr1 signalling in the development of a sexually selected trait in vertebrates, the sword of swordtail fish. BMC Dev Biol, 2008. 8: p. 98.

32. Offen, N., A. Meyer, and G. Begemann, Identification of novel genes involved in the development of the sword and gonopodium in swordtail fish. Dev Dyn, 2009. 238(7): p. 1674–87.

33. Yamamoto-Shiraishi, Y. and A. Kuroiwa, Wnt and BMP signaling cooperate with Hox in the control of Six2 expression in limb tendon precursor. Dev Biol, 2013. 377(2): p. 363–74.

34. Nachtrab, G., et al., Transcriptional components of anteroposterior positional information during zebrafish fin regeneration. Development, 2013. 140(18): p. 3754–64.

35. Moriyama, Y., et al., The medaka zic1/zic4 mutant provides molecular insights into teleost caudal fin evolution. Curr Biol, 2012. 22(7): p. 601–7.

36. Corredor-Adamez, M., et al., Genomic annotation and transcriptome analysis of the zebrafish (Danio rerio) hox complex with description of a novel member, hox b 13a. Evol Dev, 2005. 7(5): p. 362–75.

37. Waghray, A., et al., Tbx3 Controls Dppa3 Levels and Exit from Pluripotency toward Mesoderm. Stem Cell Reports, 2015. 5(1): p. 97–110.

38. Rallis, C., J. Del Buono, and M.P. Logan, Tbx3 can alter limb position along the rostrocaudal axis of the developing embryo. Development, 2005. 132(8): p. 1961–70.

39. Mise, T., et al., Function of Pax1 and Pax9 in the sclerotome of medaka fish. Genesis, 2008. 46(4): p. 185–92.

40. Amores, A., et al., A RAD-Tag Genetic Map for the Platyfish (Xiphophorus maculatus) Reveals Mechanisms of Karyotype Evolution Among Teleost Fish. Genetics, 2014. 197(2): p. 625–641.

41. Zander, C.D.D., M., Untersuchungen zur Entwicklung und Vererbung des Caudalfortsatzes der Xiphophorus-Arten(Pisces). Z Wissenschaftliche Zool 1969. 178: p. 267–315.

42. Zou, A., et al., Distribution and functional properties of human KCNH8 (Elk1) potassium channels. Am J Physiol Cell Physiol, 2003. 285(6): p. C1356–66.

43. Cloud-Richardson, K.M., B.R. Smith, and S.J. Macdonald, Genetic dissection of intraspecific variation in a male-specific sexual trait in Drosophila melanogaster. Heredity (Edinb), 2016. 117(6): p. 417–426.

44. Ichimura, K., et al., Medaka fish, Oryzias latipes, as a model for human obesity-related glomerulopathy. Biochem Biophys Res Commun, 2013. 431(4): p. 712–7.

45. Fairbairn, D., Odd Couples: Extraordinary Differences between the Sexes in the Animal Kingdom. 2013: Princeton.

46. Kallman, K.D., A new look at sex determination in Poeciliid Fishes, in Evolutionary Genetics of Fishes, B.J. Turner, Editor. 1984, Plenum Publishing Corporation. p. 95–171.

47. Schartl, M., et al., A primer of sex determination in poeciliids, in Ecology and Evolution of Poeciliid Fishes, A. Pilastro, J. Evans, and I. Schlupp, Editors. 2010. p. 264–275.

48. Powell, D.P., C.; Keegan, M.; Banerjee, S.M.; Cui, R.; Andolfatto, P., Schumer, M.; Rosenthal, G.G., The genetic architecture of the sexually selected sword ornament. BioRxiv, 2020. doi: https://doi.org/10.1101/2020.07.23.218164.

49. Perathoner, S., et al., Bioelectric signaling regulates size in zebrafish fins. PLoS Genet, 2014. 10(1): p. e1004080.

50. Lanni, J.S., et al., Integrated K+ channel and K+Cl− cotransporter functions are required for the coordination of size and proportion during development. Dev Biol, 2019. 456(2): p. 164–178.

51. Stewart, S.L.B., H.K.; Yette, G.A.; Henner, A.L.; Braunstein, J.A., Stankunas, K., longfin causes cis-ectopic expression of the kcnh2a ether-a-go-go K+ channel to autonomously prolong fin outgrowth. . BioRxiv, 2019.

52. Kon, T., et al., The Genetic Basis of Morphological Diversity in Domesticated Goldfish. Curr Biol, 2020. 30(12): p. 2260–2274 e6.

53. McLaughlin, K.A. and M. Levin, Bioelectric signaling in regeneration: Mechanisms of ionic controls of growth and form. Dev Biol, 2018. 433(2): p. 177–189.

54. Sundelacruz, S., M. Levin, and D.L. Kaplan, Role of membrane potential in the regulation of cell proliferation and differentiation. Stem Cell Rev Rep, 2009. 5(3): p. 231–46.

55. Urrego, D., et al., Potassium channels in cell cycle and cell proliferation. Philos Trans R Soc Lond B Biol Sci, 2014. 369(1638): p. 20130094.

56. Jung, H. and O. Akkus, Activation of intracellular calcium signaling in osteoblasts colocalizes with the formation of post-yield diffuse microdamage in bone matrix. Bonekey Rep, 2016. 5: p. 778.

57. Goltzman, D. and G.N. Hendy, The calcium-sensing receptor in bone--mechanistic and therapeutic insights. Nat Rev Endocrinol, 2015. 11(5): p. 298–307.

58. Han, B., et al., Eag1 K(+) Channel: Endogenous Regulation and Functions in Nervous System. Oxid Med Cell Longev, 2017. 2017: p. 7371010.

59. Somarelli, J.A., et al., Structure-based classification of 45 FK506-binding proteins. Proteins, 2008. 72(1): p. 197–208.

60. Kujawski, S., et al., Calcineurin regulates coordinated outgrowth of zebrafish regenerating fins. Dev Cell, 2014. 28(5): p. 573–87.

61. Jan, L.Y. and Y.N. Jan, Voltage-gated potassium channels and the diversity of electrical signalling. J Physiol, 2012. 590(11): p. 2591–9.

