## Supplemental Material for "The developmental and genetic architecture of the sexually selected male ornament of swordtails"

### Material and Methods

#### Experimental Animals

All fish were reared under a standard conditions [1] with a light/dark cycle of 14/10 h at 26 °C in the fish facility of the Biocenter at the University of Wuerzburg, Germany. All animals were kept and sampled in accordance with the applicable EU and national German legislation governing animal experimentation. In particular, all experimental protocols were approved through an authorization (568/300-1870/13) of the Veterinary Office of the District Government of Lower Franconia, Germany, in accordance with the German Animal Protection Law (TierSchG).

For regeneration experiments, fish were immobilized by dipping into 4°C water, and the caudal margin of the tail fin was resected with a razor blade. Tissues were collected at different stages of regeneration (fig. S2). Samples from *X. hellerii* females and the swordless males of *Priapella lacondonae* and *X.maculatus* were taken after caudal fin resection at the same day according to male sword regeneration stages. Tissues from naturally developing swords and the median and upper caudal fin margin of male *X. hellerii* were sampled at different stages according to fig. S1. Induction of the sword in mature female *X. hellerii* (4-5 months old) was done by addition of 17-methyl testosterone to the tank water (30 µg/l = 1µMol, replenished daily). The dorsal, median and ventral caudal fin margins, including the sword were collected after 11 days of treatment at a stage corresponding to naturally developing sword stage 4 (fig. S1). Areas used for RNA-seq and qPCR experiments are depicted in fig. S15. Samples from 15 – 20 individuals were pooled for RNA extraction.

#### RNA-seq transcriptomics

Total RNA was isolated using TRIzol Reagent (Thermo Fisher Scientific, Waltham, USA) according to the supplier's recommendation. Custom sequencing (BGI, Shenzen, China) of TruSeq libraries generated 25-30 million 100bp paired end reads for each sample on the Illumina Hiseq4000 platform.

#### Differential gene expression analysis

After duplicate and barcode removal reads were aligned to the Xiphophorus_hellerii-4.1 genome (https://www.ncbi.nlm.nih.gov/genome/15325?genome_assembly_id=7477339) using the STAR aligner version 2.5 (--runMode alignReads --quantMode GeneCounts) [2]. Resulting read counts were used by DESeq2 [3] for differential gene analysis. Datasets generated at different time points were analyzed separately.

For further analysis, only expressed genes were considered. “Expressed” was defined as normalized read count >= 10 in at least one sample in datasets “female” (regeneration of caudal fin in adult females), “sword development” (normal sword development in young males at puberty), testosterone induced sword in adult females (“testosterone induced sword”) or “regeneration” (regeneration of tail fin and sword in adult males). We added a published dataset (“testosterone treated juveniles”) [4] of an independent testosterone treatment for sword induction in 3 months old undifferentiated juvenile *X. hellerii*. Because dataset “testosterone treated juveniles” has four replicates for each sample a gene was required to have a normalized read count>= 10 in at least two samples. Subsequently all datasets were filtered for genes with a log2 fold change >=1 up or down, respectively, in at least one time point. Differentially expressed genes of the four male datasets were represented in a Venn diagram (https://bioinfogp.cnb.csic.es/tools/venny/) (fig. S3) and the overlap of all four datasets generated dataset “common in all male” (table S1). Next, all genes that showed the same differential regulation in “female” were removed from “common in all male”, and the remaining 54 genes (table S1) were annotated for their chromosomal location.

**qPCR expression analysis**

Total RNA was isolated from pooled samples using TRIzol Reagent (Thermo Fisher Scientific, Waltham, USA) according to the supplier’s recommendation. After DNase treatment, total RNA (1-2 µg) was reverse transcribed using the RevertAid First Strand cDNA Synthesis kit (Thermo Fisher Scientific, Waltham, USA) and random hexamer primers, according to the manufacturer's instructions. For real-time qRT-PCR, cDNA from 50 ng of total RNA was used. All results reported here are averages of at least two independent reverse transcription (RT) reactions and two PCR experiments from each such reaction. Primer sequences are listed in table S4. Amplification was monitored using a Mastercycler ep realplex$^2$ (Eppendorf, Hamburg, Germany). For quantification, expression of each gene was normalized to the housekeeping gene *ef1a1* (elongation factor 1 alpha 1) using the delta Ct method [5].

qPCR expression analysis was performed to confirm differential expression results from the RNA-seq datasets from *X. hellerii* and to monitor differential expression in other species (*X. maculatus, X. montezumae*, *X. monticolus*, *X. pygmaeus*, *P. lacandonae*, *O. latipes*) (figs. S4-10). Transcript abundance was also measured for several genes located under the chromosome 13 QTL peak to confirm that they are not differentially expressed in sword regeneration (figs. S19-20), *for tyrp1*, a marker for pigment cell differentiation (fig. 21) and *msxC* (fig. S22), a previously postulated sword gene candidate [6].

**Sequence alignment of *kcnh8***

Protein sequences of Kcnh8 were retrieved for different species: *X.hellerii* and *X.couchianus* from NCBI (XP_032437747.1, XP_027893054.1); *X.maculatus* from Ensembl (ENSXMAP00000000856); *X.birchmanni* and *X.malinche*, from a previous study [7]; *X.signum*, *X.mixei*, *X.montezumae*, *X.clemenciae*, *X.monticolus*, *X.kallmani*, *X.mayae*, *X.andersi*, *X.pygmaeus*, *X.continens*, *X.multilineatus*, *X.nigrensis*, *X.milleri*, *X.gordoni*, *X.meyeri*, *X.evelynae*, *X.xiphidium* and *X.variatus*, from raw NGS reads.

To retrieve the sequence from raw NGS reads, first, we collected all related reads by aligning them to the existing protein sequences from reference genomes using DIAMOND [8]. The kept reads were then assembled into exon-fragments using CAP3 [9]. For each fragment we determined its best translation frame by mapping it onto the reference protein sequences using GeneWise [10]. Finally, the resulting protein fragments were ordered and merged into a complete sequence according to the alignment.

**QTL mapping**

To identify regions of the genome associated with the sword trait, the Sword Index (SI), which is the sword length divided by standard length, was determined. F1 individuals were obtained from a cross of a female *Xiphophorus hellerii* (Rio Lancetilla strain) with a male *X. maculatus* (Jp163A strain) aided by artificial insemination. The low fertility of F1 intercrosses [11] precluded the production of F2 families, so we performed two backcrosses of *X. maculatus* /*X. hellerii* F1 males with *X. hellerii* (Rio Lancetilla strain) females as the recurrent parent. Quantitative trait locus (QTL) analysis was performed in R/qtl v.1.39-5 [12] with phenotype (herein) and genotype data for 85 males and 16,250 RAD-tag loci and the genetic map from Amores and colleagues [13]. Backcross generation males for mapping were produced by two sires; 60 offspring from male #2059 crossed with four full-sib females (44, 2, 12, and 2 offspring per female) and 25 from male #2074, all from one female. The dataset was coded as homozygous for the genotype of the backcross parent *X. hellerii* (data code b), or heterozygous (h) with alleles from *X. hellerii* and *X. maculatus*. Interval mapping was performed using the non-parametric model due to the non-normal distribution of the SI phenotype. Genotype probabilities were calculated at a maximum distance of 1 centiMorgan and markers with identical genotypes were removed from the analysis. The genome-wide significance threshold was determined using a permutation test with 1000 replicates. The marker sequences (table S5) used for QTL mapping were later aligned to the *X. maculatus* genome (NCBI GCF_002775205.1) and the *X. hellerii* genome (GCA_003331165.2) with GSNAP version 2018-03-25 [14] (table S3).

**Electrophysiology**

To generate cRNA for functional characterization of *Xiphophorus hellerii* Kcnh8 in *Xenopus* oocytes, the coding sequence of *Xiphophorus hellerii kcnh8* was cloned into oocyte expression vector pNBI16 (pGEM-based vector) using the USER-technique [15]. The construct was verified by sequencing. cRNA of *kcnh8* was prepared using the AmpliCap-Max™ T7 High Yield Message Maker Kit (Cellscript, Biozym Scientific GmbH, Hessisch Oldendorf, Germany). Oocyte preparation and cRNA injection have been described elsewhere [16]. Following the injection of 20 ng cRNA per oocyte, oocytes were incubated at 16°C for 24 to 36 hours in ND96 solution (96 mM NaCl, 2 mM KCl, 1 mM $CaCl_2$, 1 mM $MgCl_2$, 10 mM Hepes pH7,4) supplemented with 50 mg/l gentamycin.

In two-electrode voltage-clamp studies, oocytes were perfused with KCl-containing solutions, based on Tris/Mes buffers. The standard solution contained 10 mM Tris/Mes, pH 7.5, 1 mM $CaCl_2$, 1 mM $MgCl_2$, 30 mM KCl and 70 mM LiCl. If appropriate, osmolarity was adjusted to 220 mOsmol/kg using D-sorbitol. For measurements at varying $K^+$ concentrations, the ionic strength was kept constant by replacing KCl with LiCl and vice versa. Voltage-dependent activation of Kcnh8-expressing oocytes was recorded with voltage-pulse-protocols designed and applied with the acquisition software Patchmaster (HEKA Elektronik GmbH, Lambrecht/Pfalz, Germany). Proceeding from a holding potential ($V_H$) of -20 mV, a series of 4s test voltage pulses ranging from +40 to -140 mV in 10 mV decrements were applied. Steady state currents ($I_{SS}$) were extracted at the end of the test voltage pulses.

### References


1. Kallman, K., *The platyfish, Xiphophorus maculatus*, in *Handbook of Genetics*, K. RC, Editor. 1975, Plenum Press: New York, N.Y. p. 81-132.
2. Dobin, A., et al., *STAR: ultrafast universal RNA-seq aligner.* Bioinformatics, 2013. **29**(1): p. 15-21.
3. Love, M.I., W. Huber, and S. Anders, *Moderated estimation of fold change and dispersion for RNA-seq data with DESeq2.* Genome Biol, 2014. **15**(12): p. 550.
4. Kang, J.H., et al., *Transcriptomics of two evolutionary novelties: how to make a sperm-transfer organ out of an anal fin and a sexually selected "sword" out of a caudal fin.* Ecol Evol, 2015. **5**(4): p. 848-64.
5. Simpson, D.A., et al., *Retinal VEGF mRNA measured by SYBR green I fluorescence: A versatile approach to quantitative PCR.* Mol Vis, 2000. **6**: p. 178-83.
6. Zauner, H., et al., *Differential regulation of msx genes in the development of the gonopodium, an intromittent organ, and of the "sword," a sexually selected trait of swordtail fishes (Xiphophorus).* Evol Dev, 2003. **5**(5): p. 466-77.
7. Powell, D.L., et al., *Natural hybridization reveals incompatible alleles that cause melanoma in swordtail fish.* Science, 2020. **368**(6492): p. 731-736.
8. Buchfink, B., C. Xie, and D.H. Huson, *Fast and sensitive protein alignment using DIAMOND.* Nature methods, 2015. **12**(1): p. 59-60.
9. Huang, X. and A. Madan, *CAP3: A DNA sequence assembly program.* Genome research, 1999. **9**(9): p. 868-877.

10. Birney, E., M. Clamp, and R. Durbin, *GeneWise and genomewise.* Genome research, 2004. **14**(5): p. 988-995.
11. Franchini, P., et al., *Long-term experimental hybridisation results in the evolution of a new sex chromosome in swordtail fish.* Nat Commun, 2018. **9**(1): p. 5136.
12. Broman, K.W., et al., *R/qtl: QTL mapping in experimental crosses.* Bioinformatics, 2003. **19**(7): p. 889-90.
13. Amores, A., et al., *A RAD-Tag Genetic Map for the Platyfish (Xiphophorus maculatus) Reveals Mechanisms of Karyotype Evolution Among Teleost Fish.* Genetics, 2014. **197**(2): p. 625-641.
14. Wu, T.D. and C.K. Watanabe, *GMAP: a genomic mapping and alignment program for mRNA and EST sequences.* Bioinformatics, 2005. **21**(9): p. 1859-75.
15. Nour-Eldin, H.H., et al., *Advancing uracil-excision based cloning towards an ideal technique for cloning PCR fragments.* Nucleic Acids Res, 2006. **34**(18): p. e122.
16. Becker, D., et al., *Changes in voltage activation, Cs+ sensitivity, and ion permeability in H5 mutants of the plant K+ channel KAT1.* Proc Natl Acad Sci U S A, 1996. **93**(15): p. 8123-8.

### Supplementary Figures

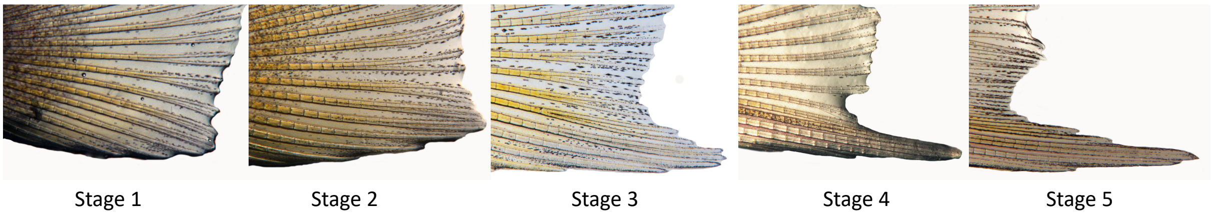


**Fig. S1: Stages of normal sword development in *Xiphohorus he*llerii males during puberty.**

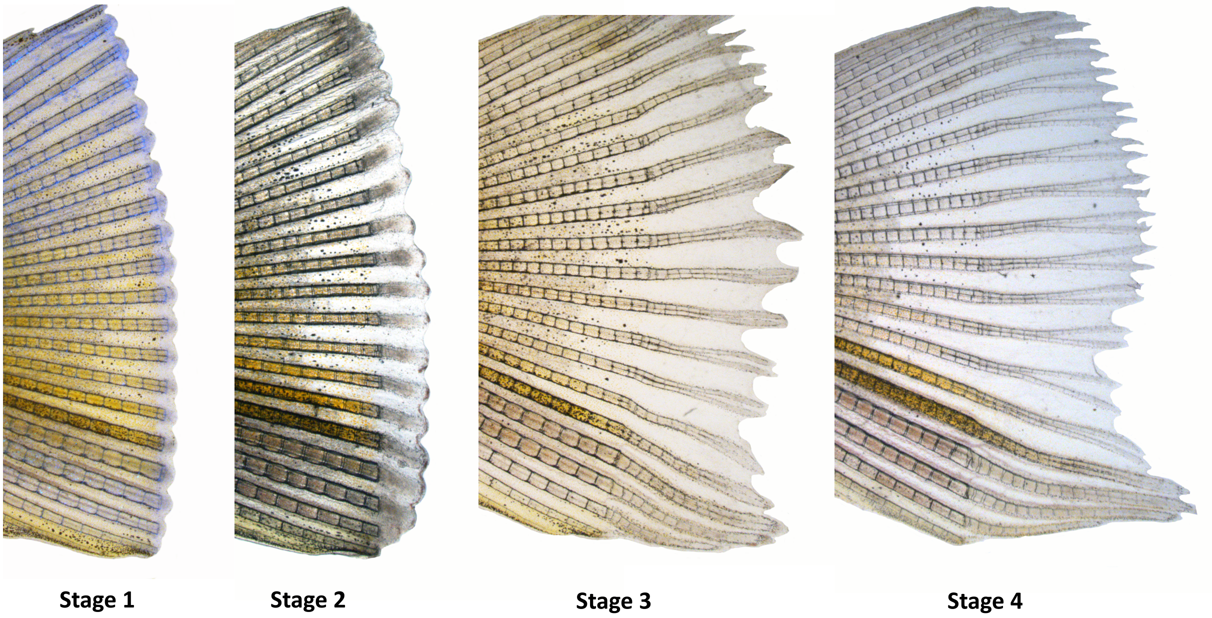


**Fig. S2: Stages of sword regeneration in *Xiphohorus hellerii* males.**

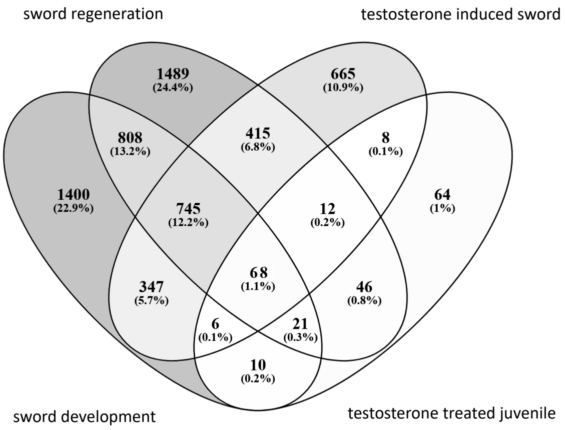


**Fig. S3. Venn diagram of differentially expressed genes.** Numbers of genes with log2FC>=1 between upper and lower caudal fin margin during natural sword development (stage 1-5), sword regeneration (days 0-10), testosterone induced sword in females and testosterone treated juvenile *Xiphophorus hellerii*.

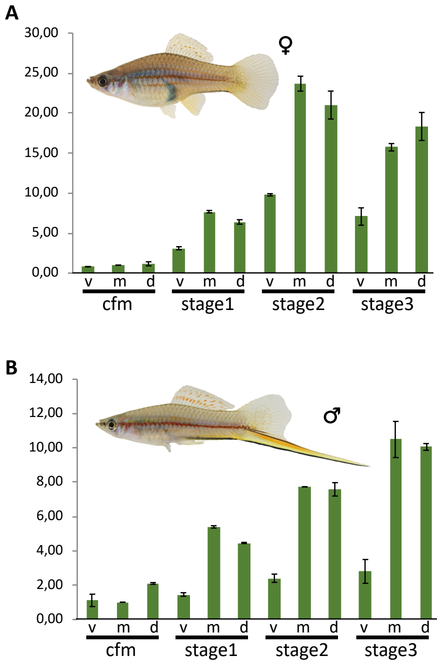


**Fig. S4: Establishment of a spatial expression pattern of leukocyte receptor kinase (*ltk*) in the caudal fin during regeneration.** Expression of ltk in the caudal fin margin of the tail (cfm) and during regeneration stages of adult *Xiphophorus hellerii* females (**A**) and males (**B**) (v, ventral, m, median, d, dorsal compartment). Vertical axis indicates fold change of expression normalized to cfm, m.

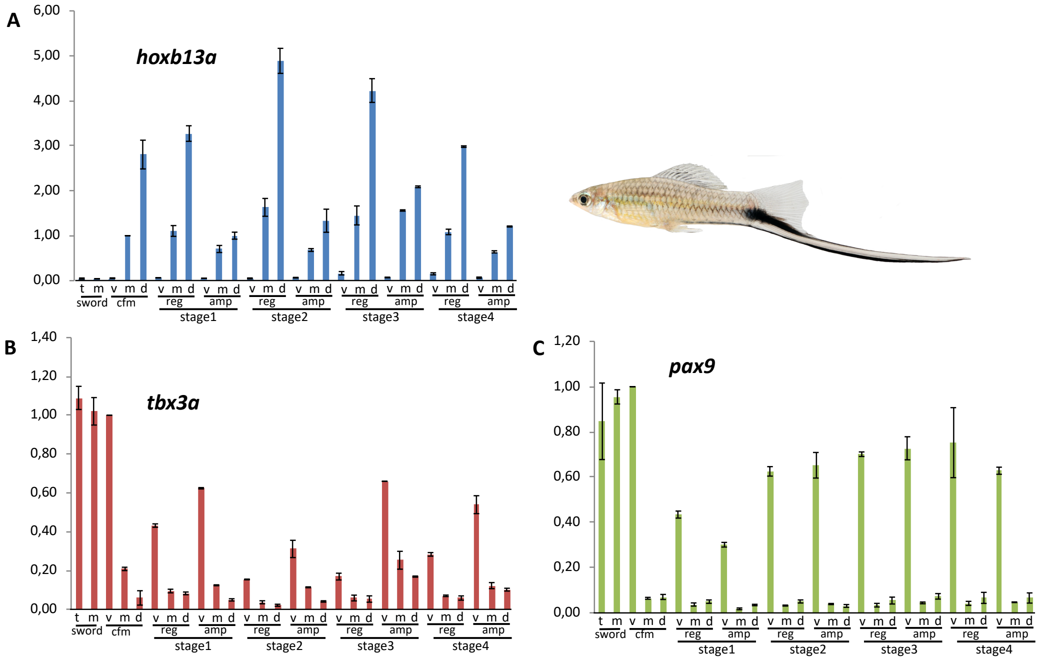


**Fig. S5: Spatial expression pattern of transcription factor genes in the caudal fin and sword of male *Xiphophorus montezumae*.** Expression of *hoxb13a* (**A**)*, tbx3a* (**B**) and *pax9* (**C**) in the caudal fin margin of the tail fin (cfm), the median sector (m) and tip (t) of the sword and during sword regeneration (v, ventral, m, median, d, dorsal compartment) in the regenerating tissue (reg) and the compartment proximal to the regenerate (amp). Vertical axis indicates fold change of expression normalized to cfm, v (*tbx3a, pax9*) or cvm, m (*hoxb13a*).

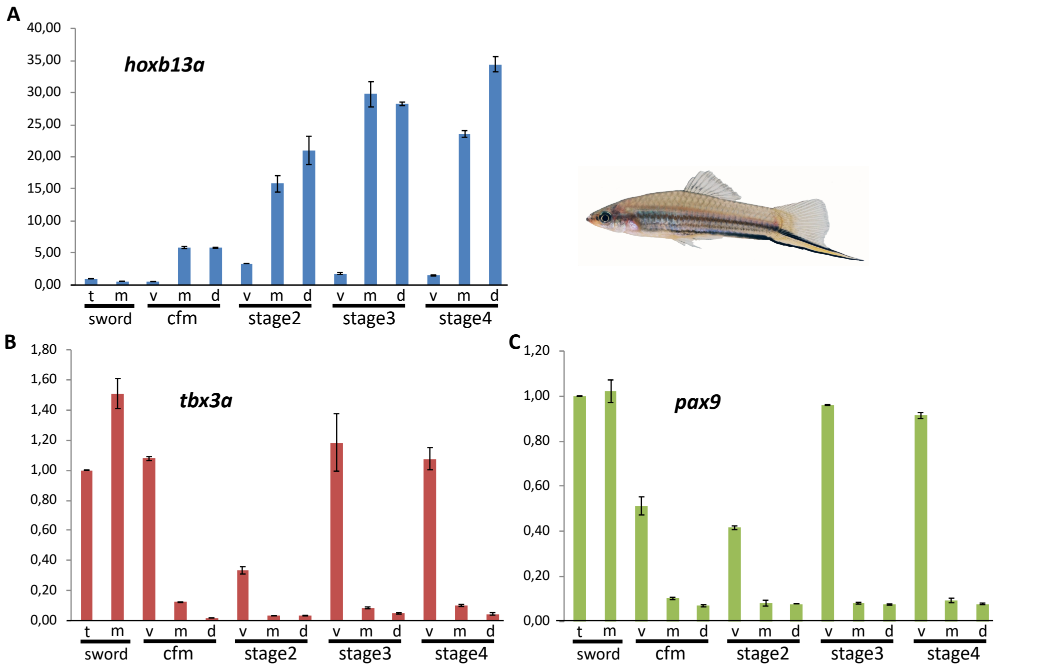


**Fig. S 6: Spatial expression pattern of transcription factor genes in the caudal fin and sword of male *Xiphophorus monticolus*.** Expression of transcription factor genes *hoxb13a* (**A**), *tbx3a* (**B**) and *pax9* (**C**) in the caudal fin margin of the tail fin (cfm) of adult *Xiphophorus monticolus* males, the median sector (m) and tip (t) of the sword and during sword regeneration (v, ventral, m, median, d, dorsal compartment). Vertical axis indicates fold change of expression normalized to sword, t.

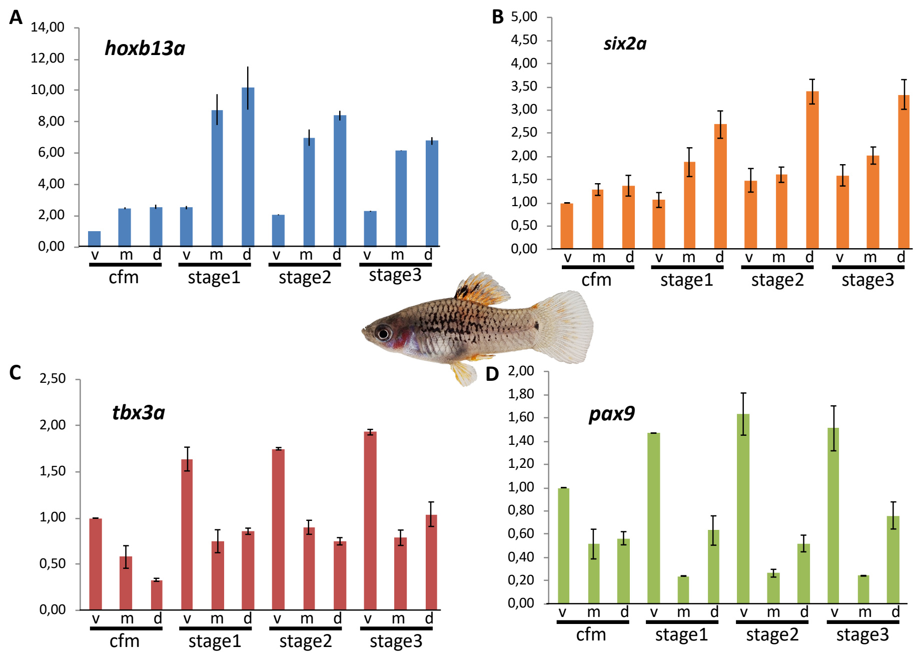


**Fig. S7: Spatial expression pattern of transcription factor genes in the caudal fin of male *Xiphophorus maculatus*.** Expression of transcription factor genes *hoxb13a*, *six2a*, *tbx3a* and *pax9* in the caudal fin margin of the tail fin (cfm) and during tail fin regeneration (v, ventral, m, median, d, dorsal compartment). Vertical axis indicates fold change of expression normalized to cfm, v.

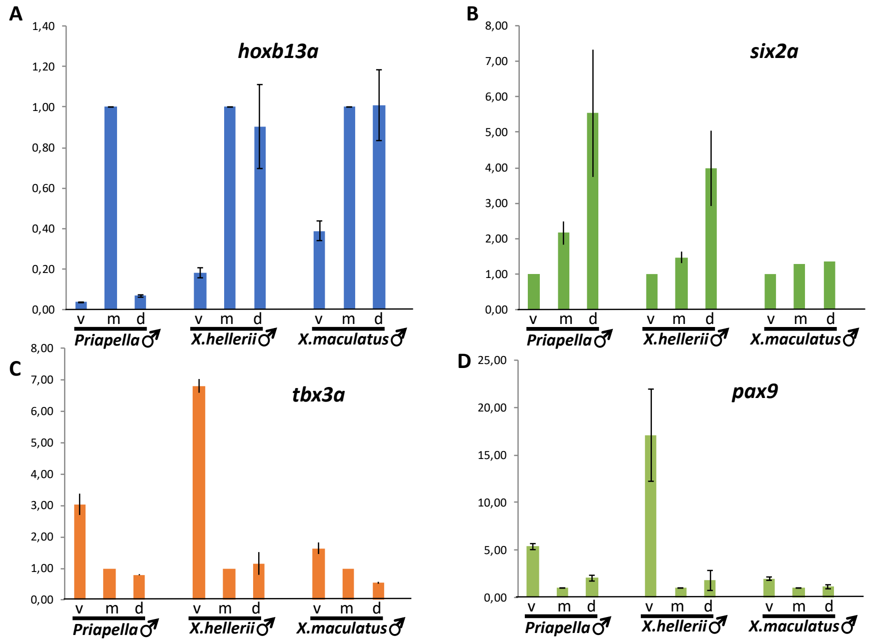


**Fig. S8: Comparison of transcription factor expression patterns in different species.** Expression of transcription factor genes *hoxb13a* (**A**), *six2a* (**B**), *tbx3a* (**C**) and *pax9* (**D**) in the caudal fin margin of the tail fin of adult males of *Priapella lacandonae*, *Xiphophorus hellerii* and *Xiphophorus maculatus*. (v, ventral, m, median, d, dorsal compartment). Vertical axis indicates fold change of expression normalized to cfm, m (A,C,D) or cfm, v(B).

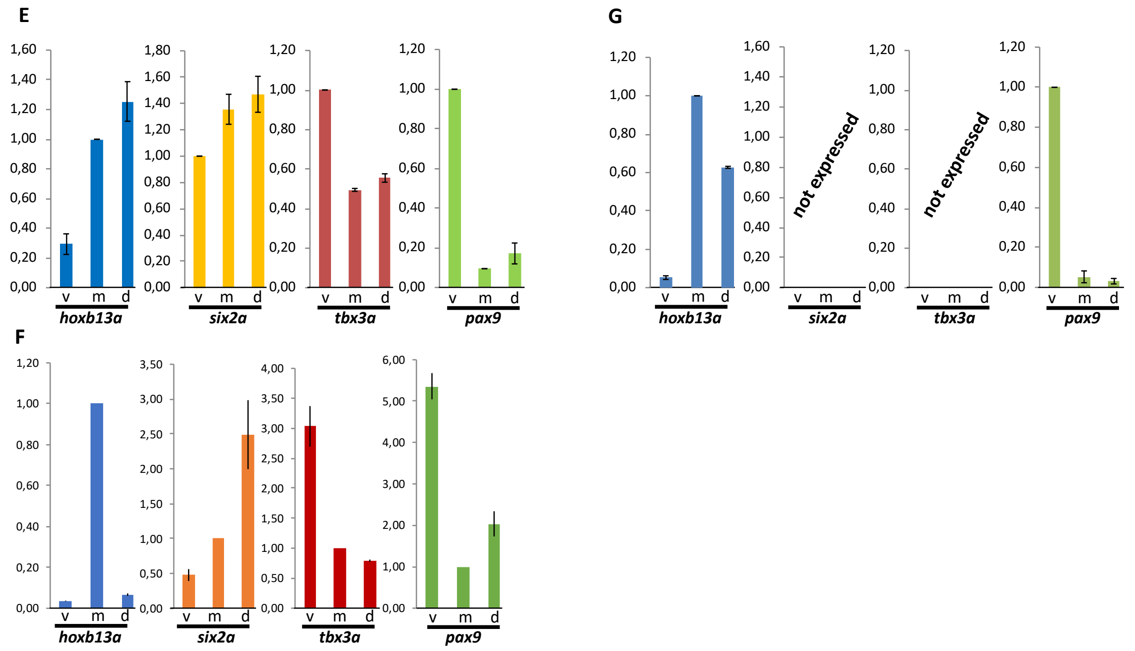


**Fig. S9: Comparison of transcription factor expression patterns in different species.** Expression of *hoxb13a*, *six2a*, *tbx3a* and *pax9* in the caudal fin margin of the tail fin (cfm) of (**E**) adult pygmy swordtails, *Xiphophorus pygmaeus*, (**F**) *Priapella lacandonae* and (**G**) medaka, *Oryzias latipes.* v, ventral, m, median, d, dorsal compartment. Vertical axis indicates fold change of expression normalized to cfm, m, except for medaka *pax9* and *X. pygmaeus six2a*, *tbx3a* and *pax9*, cfm, v.

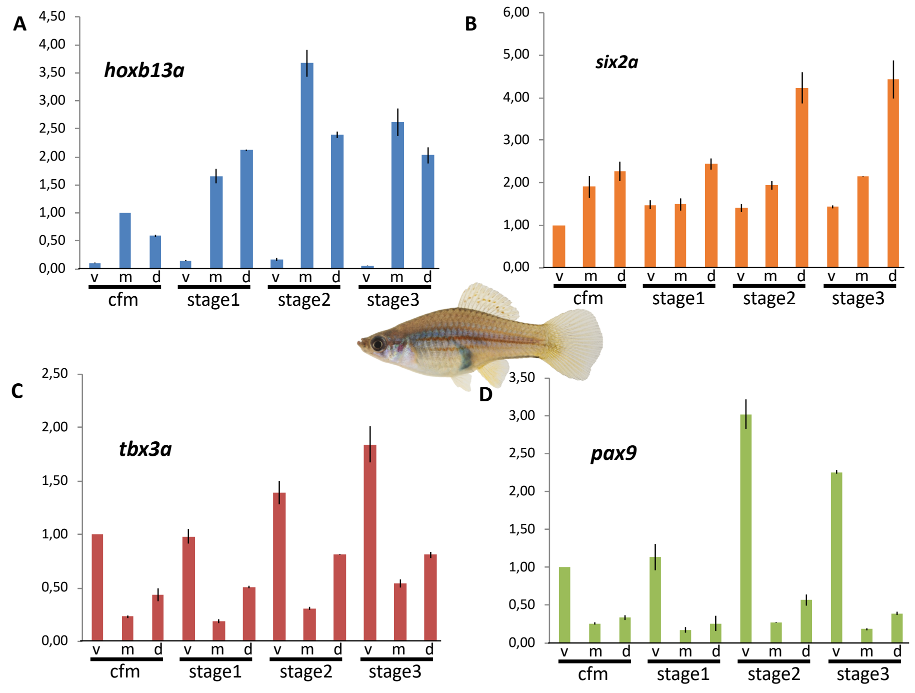


**Fig. S10: Spatial expression pattern of transcription factor genes in the caudal fin of *Xiphophorus hellerii* females.** Expression of *hoxb13a* (**A**), *six2a* (**B**), *tbx3a* (**C**) and *pax9* (**D**) in the caudal fin margin of the tail fin (cfm) and during tail fin regeneration (v, ventral, m, median, d, dorsal compartment). Vertical axis indicates fold change of expression normalized to cvm, v (*six2a, tbx3a, pax9*) or cfm, m (*hoxb13a*).

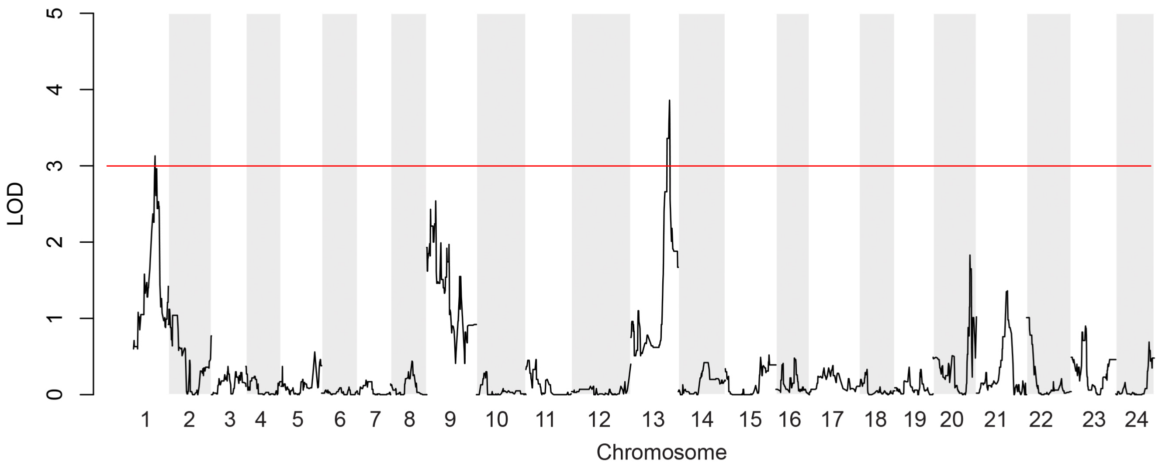


**Fig. S11. Manhattan plot of quantitative trait loci (QTL) mapping results for sword length**. One major QTL peak is located on chromosome 13, two minor peaks on chromosomes 1 and 9, and several smaller peaks on chromosomes 20 – 24. The plot depicts aligned RAD-tag positions on the *Xiphophorus hellerii* genome version 4.1 with maximum likelihood statistics.

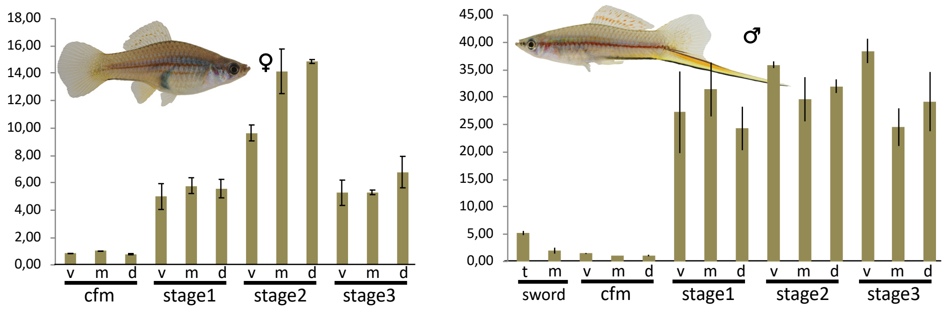


**Fig. S12: Spatial expression pattern of *fkbp9*.** Expression in the caudal fin margin of the tail fin (cfm) of adult *Xiphophorus hellerii* females (left) and males (right), the median sector (m) and tip (t) of the sword and during regeneration (v, ventral, m, median, d, dorsal compartment). Vertical axis indicates fold change of expression normalized to cfm, m.

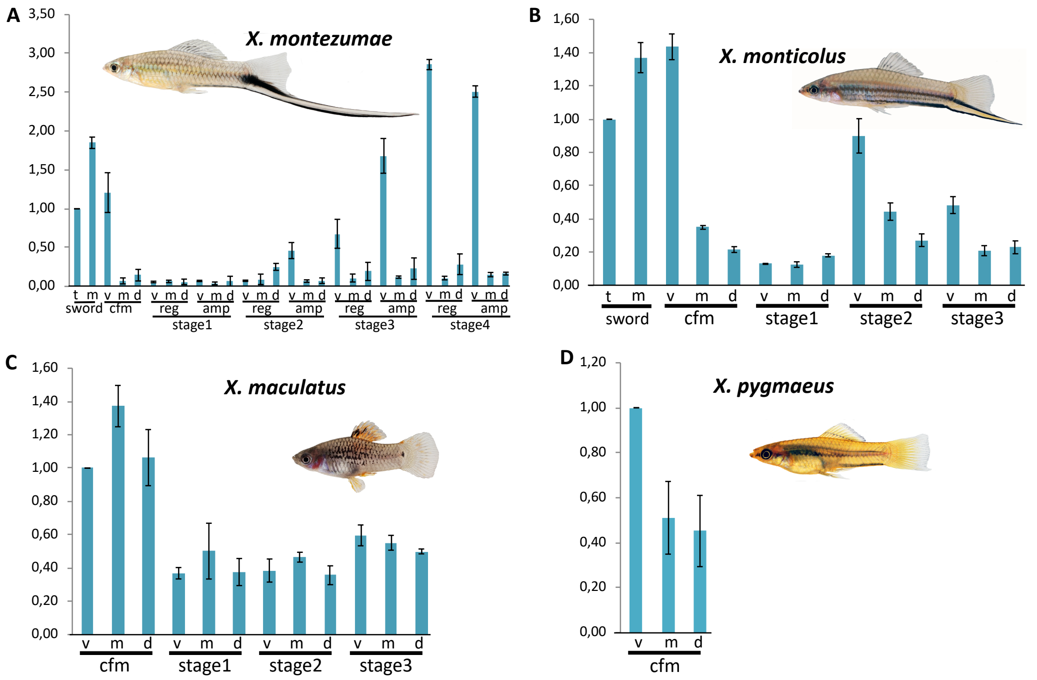


**Fig. S13: Spatial expression pattern of transcription factor genes in the caudal fin of *Xiphophorus* species.** Expression of *kcnh8* in the caudal fin margin of the tail fin (cfm) of adult *Xiphophorus montezumae* (**A**)*, X. monticolus* (**B**)*, X. maculatus* (**C**) and *X. pygmaeus* (**D**) males, the median sector (m) and tip (t) of the sword and during sword regeneration (v, ventral, m, median, d, dorsal compartment). Vertical axis indicates fold change of expression normalized to sword, t (A), (B) and cfm, v (C), (D).

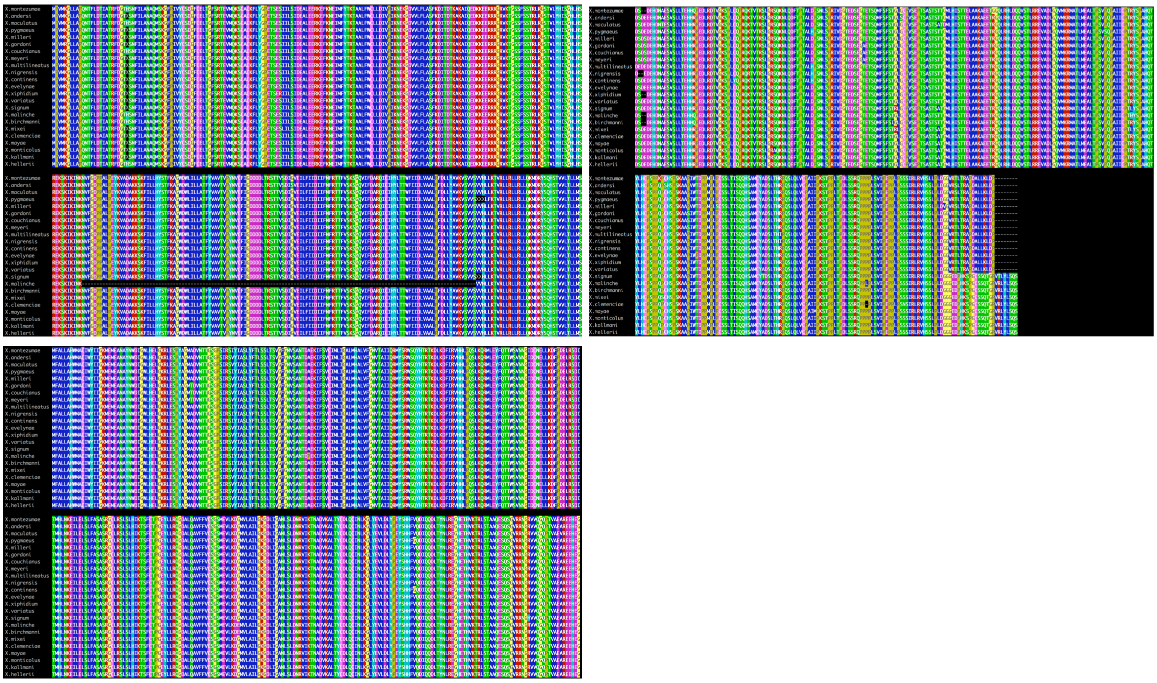

**Fig. S14: Alignment of protein sequences of Kcnh8 from *Xiphophorus* species.** The missing sequence from X. malinche (corresponding to one exon) is most likely due to a misassembly.

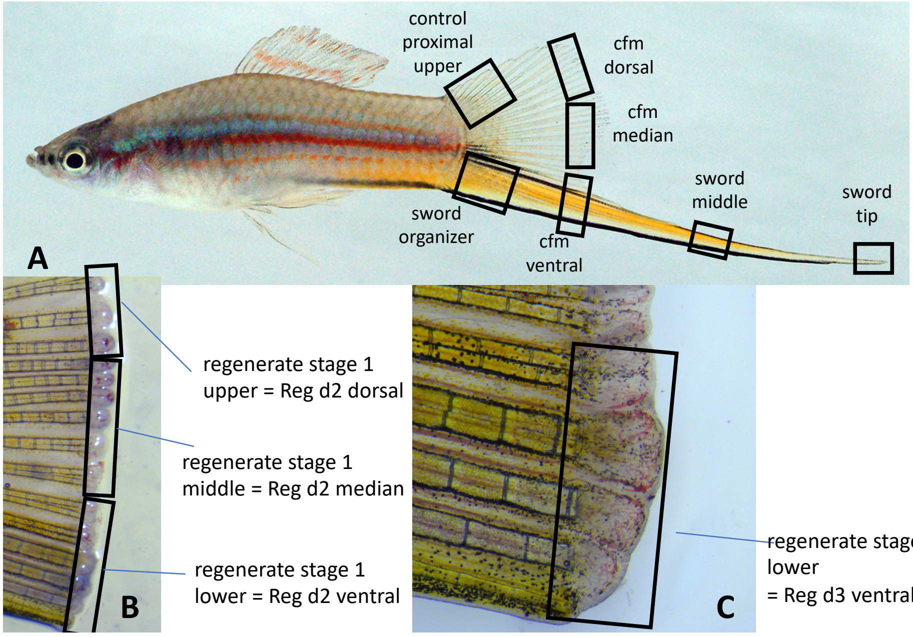


**Fig. S15**: **Compartments used for sampling** (indicated by black boxes). (**A**) Regions of the tail fin taken for amputation. Cfm. Caudal fin margin (**B**) Regenerate blastema at 2 dpa and regions taken for RNA extractions (**C**) Regenerate blastema at 3dpa, the ventral part (boxed) starts to grow more than median and dorsal

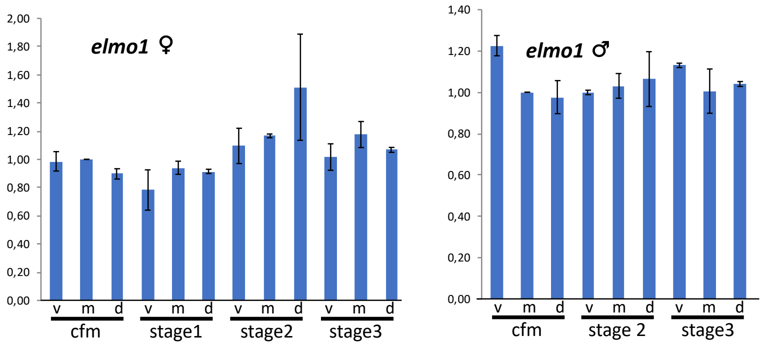


**Fig. S16: Expression pattern of *elmo1*, a gene located on chromosome 13 in the QTL region.** Expression of *elmo 1*in the caudal fin margin of the tail fin (cfm) of adult *Xiphophorus hellerii* females (left) and males (right) Vertical axis indicates fold change of expression normalized to cfm, m.

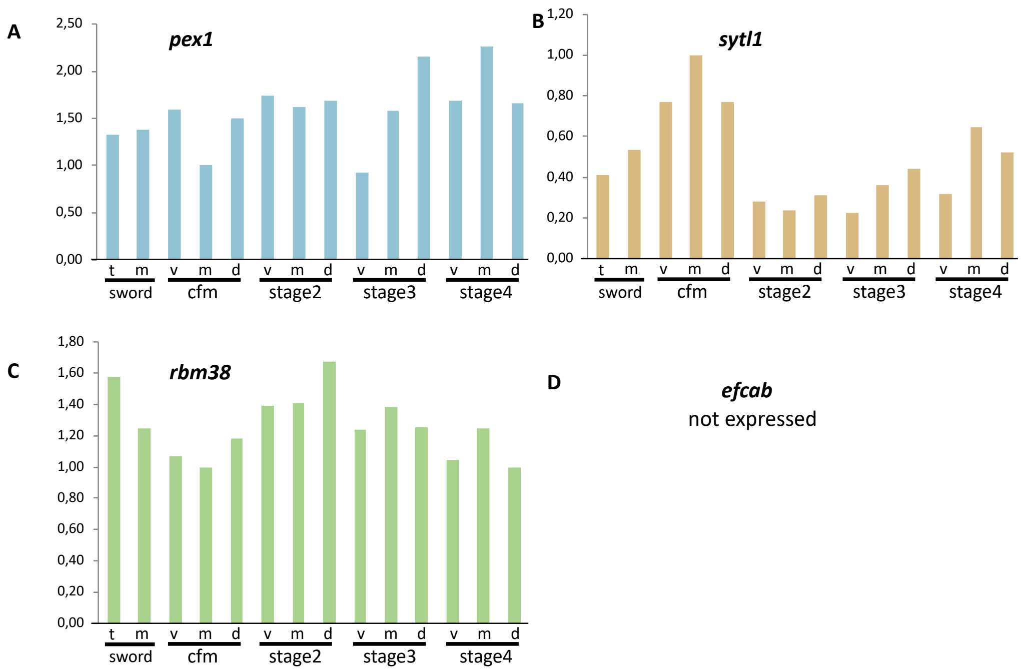


**Fig. S17: Expression pattern of genes located on chromosome 13 in the QTL region, in the caudal fin and sword of male *Xiphophorus hellerii*** Expression of *pex1* (**A**), *sytl1* (**B**) and *rbm38* (**C**) in the caudal fin margin of the tail fin (cfm), the median sector (m) and tip (t) of the sword and during sword regeneration (v, ventral, m, median, d, dorsal compartment). *efcab* expression (**D**) was not detected. Vertical axis indicates fold change of expression normalized to cfm, m.

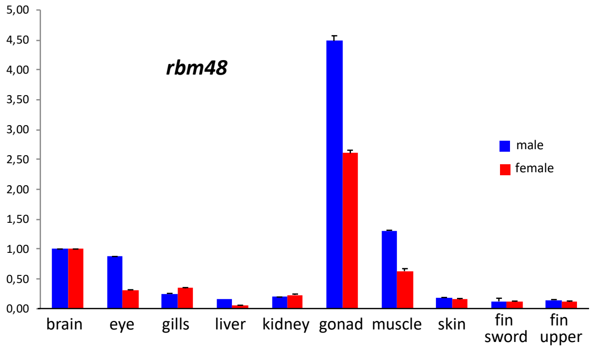


**Fig. S18: Expression pattern of *rbm48*, a gene located on chromosome 13 in the QTL region in *Xiphophorus hellerii*.** No differential expression between males and females in the caudal fin was detected. Vertical axis indicates fold change of expression normalized to brain (A),

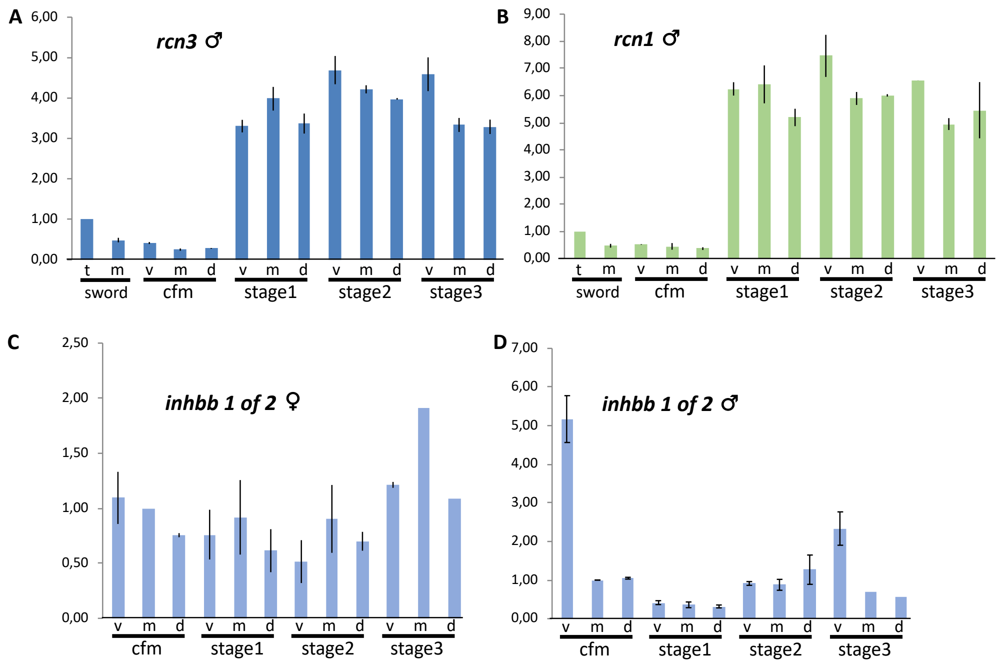


**Fig. S19: Expression patterns of genes that are regulated during regeneration.** Expression of *rcn3* (**A**) and *1* (**B**) and *inhbb1 of 2* (**C,D**) in the caudal fin margin of the tail fin (cfm) (v, ventral, m, median, d, dorsal compartment) of male (A,B,D) and female (C) *Xiphohorus hellerii*. Vertical axis indicates fold change of expression normalized to cfm, v (A,B) or cfm, m (C,D).

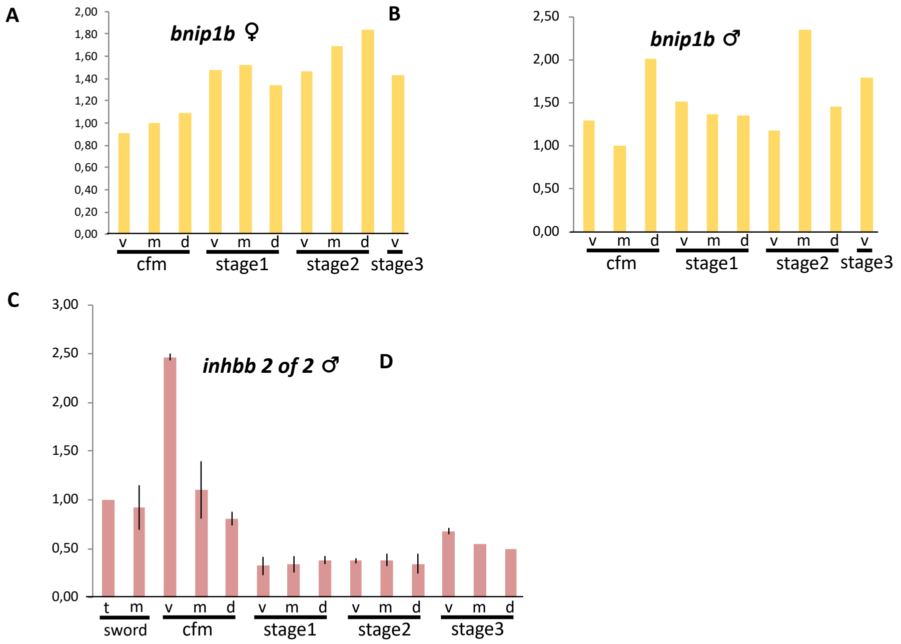


**Fig. S20: Expression patterns of genes that are regulated during regeneration.** Expression of *bnip1b* (**A, B**) and *inhbb2 of 2* (**C**) in the caudal fin margin of the tail fin (cfm) (v, ventral, m, median, d, dorsal compartment) of male (B, C) and female (A) *Xiphohorus hellerii* and during regeneration stages. Vertical axis indicates fold change of expression normalized to cfm, m (A, B) or sword, t.

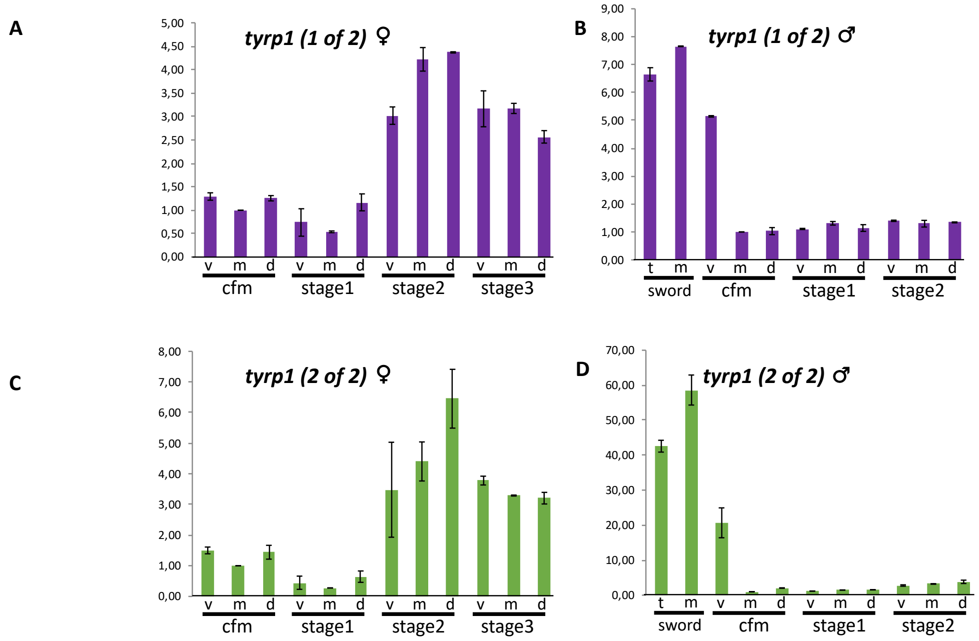


**Fig. S21: Expression of the pigmentation gene *tyrosinase related protein 1* (*tyrp1*).** Expression of *tyrp1* ohnologs in the caudal fin margin of the tail fin (cfm) (v, ventral, m, median, d, dorsal compartment) and during regeneration stages of adult *Xiphophorus hellerii* females (A, C) and the sword in males (B, D). Vertical axis indicates fold change of expression normalized to cfm, m.

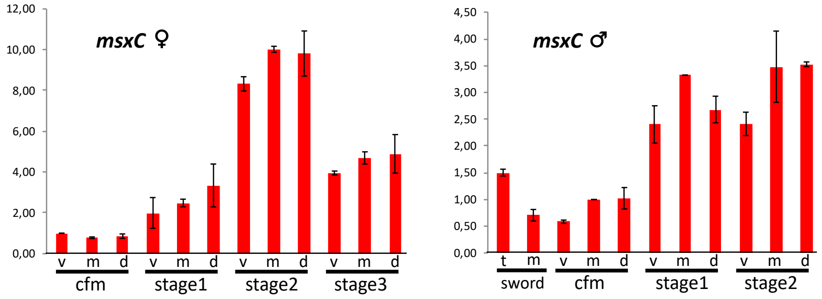


**Fig. S22: Expression of candidate gene msxC.** Expression of *msxC* in the caudal fin margin of the tail fin (cfm) (v, ventral, m, median, d, dorsal compartment) and during regeneration stages of adult *Xiphophorus hellerii* females (left) and the sword in males (right). Vertical axis indicates fold change of expression normalized to cfm, v (females) or cfm, m (males).
